# Clustering FunFams using sequence embeddings improves EC purity

**DOI:** 10.1101/2021.01.21.427551

**Authors:** Maria Littmann, Nicola Bordin, Michael Heinzinger, Konstantin Schütze, Christian Dallago, Christine Orengo, Burkhard Rost

## Abstract

**Motivation:** Classifying proteins into functional families can improve our understanding of protein function and can allow transferring annotations within one family. For this, functional families need to be “pure”, i.e., contain only proteins with identical function. Functional Families (FunFams) cluster proteins within CATH superfamilies into such groups of proteins sharing function. 11% of all FunFams (22,830 of 203,639) contain EC annotations and of those, 7% (1,526 of 22,830) have inconsistent functional annotations.

**Results:** We propose an approach to further cluster FunFams into functionally more consistent sub-families by encoding their sequences through embeddings. These embeddings originate from language models transferring knowledge gained from predicting missing amino acids in a sequence (ProtBERT) and have been further optimized to distinguish between proteins belonging to the same or a different CATH superfamily (PB-Tucker). Using distances between embeddings and DBSCAN to cluster FunFams and identify outliers, doubled the number of pure clusters per FunFam compared to random clustering. Our approach was not limited to FunFams but also succeeded on families created using sequence similarity alone. Complementing EC annotations, we observed similar results for binding annotations. Thus, we expect an increased purity also for other aspects of function. Our results can help generating FunFams; the resulting clusters with improved functional consistency allow more reliable inference of annotations. We expect this approach to succeed equally for any other grouping of proteins by their phenotypes.

**Availability:** Code and embeddings are available via GitHub: https://github.com/Rostlab/FunFamsClustering

## Introduction

Knowledge about the function of a protein is crucial for a wide array of biomedical applications. Classifying protein sequences into functional families can shed light on uncharacterized proteins. Functional families can also reveal insights into the evolution of function through sequence changes [1]. To gain meaningful insights, those families should be consistent, i.e., only contain functionally similar proteins.

CATH FunFams [2, 3] provide a functional sub-classification of CATH superfamilies [4, 5]. Superfamilies are the last level (H) in the CATH hierarchy; they group sequences which are related by evolution, often loosely referred to as homologous. While proteins in one superfamily can still be functionally and structurally diverse, *Functional Families* (FunFams) further sub-classify superfamilies into coherent subsets of proteins with the same function. FunFams can be used to predict function on a per-protein level as described through Gene Ontology (GO) terms [6, 7], to predict functional sites [8], or to improve binding residue predictions through consensus [9].

The Enzyme Commission number (EC number) [10] numerically classifies enzymatic functions based on the reactions they catalyze. It consists of four levels and each level provides a more specific description of function than the previous one. The function of two proteins is more similar, the more levels of their two EC numbers are identical, particularly, for the levels EC3 and EC4 which describe the chemical reaction and its substrate specificity.

For 22,830 FunFams (11% of all), manually curated annotations from UniProt [11] for EC numbers for all four levels are available at least for one member. By design, proteins from the same FunFam should share the same EC class (annotated up to level 4). However, 1,526 FunFams (7% of 22,830) accounting for 16% of all sequences in the 22,830 FunFams with EC annotations have more than one annotation, and 180 (1% of 22,830) accounting for 2% of the sequences even have four or more different annotations (Fig. S1 in Supporting Online Material (SOM)). Different EC annotations within one FunFam could originate from multifunctional enzymes (e.g., promiscuous enzymes [12] or moonlighting enzymes [13]). Assuming the multifunctional enzyme to have two EC numbers, only one of those could be inconsistent with the other FunFam members rendering that FunFam seemingly inconsistent. However, different EC annotations can also result from impurity, i.e., FunFams containing proteins with different functions. Splitting FunFams further could provide a more fine-grained and consistent set of functionally related proteins.

Over the last few years, novel representations (embeddings) for proteins have emerged from adapting language models (LMs) developed for natural language processing (NLP) to protein sequences [14–18]. These embeddings are learnt solely from protein sequences either through auto-regressive pre-training (predicting the next amino acid in a sequence, e.g., ELMo [19] or GPT [20]) or through masked language modeling (reconstructing corrupted amino acids from the sequence, e.g., BERT [21]) without using any annotations (self-supervised) or any knowledge of evolutionary constraints. To accomplish this, the protein LM learns some aspects of the language of life as written in protein sequences [16]. Features learnt implicitly by these models can be transferred to any task by extracting the hidden states of the LM for a given protein sequence (transfer learning). These hidden states are referred to as the embeddings of the corresponding LM. Such embeddings capture higher-level features of proteins, including various aspects of protein function [22–28]. Therefore, we hypothesized that this orthogonal perspective – using embedding rather than sequence space – might help to find functionally consistent sub-groups within protein families built using sequence similarity.

Here, we proposed a clustering approach to identify clusters in FunFams that are more consistent in terms of shared functionality. To this end, shared functionality was defined as sharing the same EC annotation up to the fourth level (i.e., completely identical EC numbers). We represented protein sequences as embeddings, i.e., fixed-size vectors derived from pre-trained LMs.

We used the LM ProtBERT [15] to retrieve the initial embeddings, and applied contrastive learning [29–31] using the triplet loss [32] to learn a new embedding space which was optimized towards maximizing the Euclidean distance between proteins from different CATH classes while minimizing the distance between proteins in the same CATH class. During training, the distance in embedding space between similar proteins is decreased and between dissimilar proteins is increased. Similarity was defined based on the CATH hierarchy.

The resulting embeddings are called PB-Tucker. Clustering was then performed based on the Euclidean distances between those embeddings using DBSCAN [33]. Within each FunFam, DBSCAN identified clusters as dense regions in which all sequences were close to each other in embedding space; it classified proteins as outliers if they were not close to other sequences in the FunFam. That allowed the identification of (i) a more fine-grained clustering of the FunFams, and (ii) single sequences which might have been falsely assigned to this FunFam. Analyzing whether or not embedding-based clustering reduced the number of different EC annotations in a FunFam allowed validating our new approach.

## Methods

### FunFams dataset

The current version of CATH (v4.3) holds 4,328 superfamilies split into 212,872 FunFams. The FunFams generation process, albeit changing through time, consists of various steps, starting with the clustering of all sequences within a CATH superfamily at 90% sequence similarity, encoding these clusters in Hidden Markov Models and creating a relationship tree between all clusters using GeMMA [34] and HHsuite [35]. Subsequently, CATH-FunFHMMer [7] is applied to traverse the tree and GroupSim [36] conservation patterns are employed to merge or cut the tree branches to obtain the largest possible, functionally pure family. CATH FunFams have higher functional purity than CATH superfamilies and conserved residues are enriched in functional sites [7].

### EC annotations and EC purity

EC annotations for the FunFams dataset were obtained using the UniProt [11] SPARQL API and cross-assigned to all UniProt IDs available within the FunFams. UniProt provides manually curated annotations combining multiple sources through UniRule [37] and using the standardized vocabulary from Rhea [38].

Since proteins in the same FunFam are assumed to share a function, we expect all proteins in one FunFam to have the same EC number(s). If not, the FunFam is considered *impure*, i.e., it contains sequences which belong in another FunFam. If all proteins were annotated with a single EC number, impurity could naively be defined as any FunFam with more than one unique EC number (i.e., at least one protein is annotated to another EC than the other family members). However, some proteins are annotated with multiple EC numbers. These proteins might either execute multiple functions [12, 13] or might be mis-classified. In fact, 8% (7,586) of all proteins with EC annotations in our final dataset are annotated to more than one EC (Table S1).

Consider the following two FunFams: FF1 has several proteins all annotated with the same two EC numbers EC1 and EC2, while FF2 has one protein with EC1 another with EC2. Then we consider FF1 as pure and FF2 as impure. The purity of clusters was defined accordingly. We only considered EC annotations with all four levels; all others were considered un-annotated.

### Embeddings representing proteins (PB-Tucker)

We used ProtBERT [15] to create fixed-length vector representations (embeddings), i.e., vectors with the same number of dimensions irrespective of protein length. ProtBERT uses the architecture of the LM BERT [21] which applies a stack of self-attention [39] layers for masked language modeling (details in Supporting Online Material Section SOM_1.2). Fixed-length vectors were derived by averaging over the representations of each amino acid extracted from its last layer. This simple global average pooling provides an effective baseline [15, 16, 23]. In the following, ProtBERT refers to this representation.

We applied contrastive learning [29–31] on the ProtBERT embeddings to learn a new embedding space which was optimized to increase the distance between CATH superfamilies and brings those within one superfamily closer while pushing members of different superfamilies apart. Towards this end, we used the triplet loss [32] to optimize Euclidean distances between protein triplets, i.e., an anchor protein is compared to a positive and a negative protein. During training, the distance in embedding space between anchor and positive is decreased, that between anchor and negative is increased. The notion of positive and negative was taken from the CATH hierarchy.

While supervised learning for a CATH-like hierarchy is challenging, using a hierarchy to define relative similarity between triplets is straightforward as anchor and positive only need to share one level more in the hierarchy than anchor and negative. Toward this end, ProtBERT representations were projected in two steps from 1024-dimensions (1024-d) to 128-d using CATH v4.3 [3] for training the two-layer neural network (SOM_1.2). In the following, we call these new 128-d embeddings PB-Tucker (Heinzinger et al., unpublished). PB-Tucker has been trained to differentiate CATH superfamilies and seemed to better capture functional relationships between proteins in one superfamily than the original ProtBERT (SOM_1.2).

### Clustering

Representing sequences as PB-Tucker embeddings, we calculated the Euclidean distance between all embeddings within one FunFam. The distance *d* between two embeddings *x* and *y* was defined as:

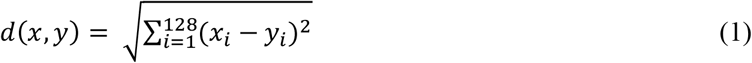

Alternatively, we tested Cosine distance (Eqn. 2), commonly used for embeddings, and Manhattan distance (Eqn. 3), commonly used for high-dimensional data.

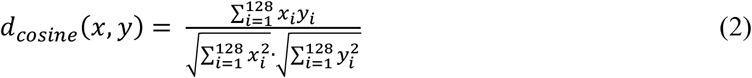

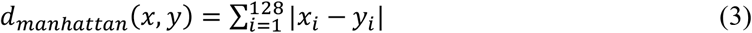

Based on the Euclidean distances, we clustered all sequences within one FunFam using the implementation of DBSCAN [33] in scikit-learn [40]. For a set of data points, DBSCAN identifies dense regions, i.e., regions of points close to each other, and classifies these regions as clusters. Data points not close to enough other data points are classified as outliers. DBSCAN identifies *core points* to seed a cluster; all points within a certain distance of the core point are added to this cluster. The two free parameters are: (1) the distance cutoff θ to consider points close (points A and B are considered close, if d(A,B)< θ), and (2) the number of neighbors *n* (including the point itself) required for a point to become “core point”; *n* implicitly controls the size and number of clusters. For our application, DBSCAN has two major advantages: (1) The number of clusters does not have to be set *a priori*, and (2) clustering and outlier detection are simultaneous.

If not stated otherwise, we used the default *n*=5 although it has been suggested to use values between *n*=D+1 and *n*=2*D−1 where D is the number of dimensions [41]. With d=128 for the PB-Tucker embeddings that implies *n*=255. Since FunFams vary in size, *n* might be adjusted to that size. For five superfamilies, we tested, in addition to *n*=5, *n*=129, *n*=255 as fixed neighborhood sizes, as well as *n*=0.05*|F|, *n*=0.1*|F|, *n*=0.2*|F| (|F|=number of sequences in FunFam) as variable neighborhood sizes dependent of the size of the FunFam.

Observing differences in the distances between the members of different superfamilies (Fig. S3), it appeared best to choose superfamily-specific values for θ. Initially, we considered using expected distance between any two members of the same FunFam. However, large distances between members in one FunFam might reveal impurity rather than a generic width of a family. Instead, we computed the median over those distances for all FunFams in one superfamily and used this value for each FunFam. The resulting value still reflects the expected distance between pairs, but the effect of large distances due to impurity should be averaged out by considering all FunFams in a superfamily. In detail, for each member in each FunFam, we calculated its average distance to all other members of that FunFam (distance distribution for five superfamilies in Fig. S3). Given the distribution of these average sequence distances, we chose the median distance as θ, i.e., we chose a distance cutoff so that 50% of all sequences in a superfamily were on average within a distance of θ to all other sequences in the same FunFam. Decreasing θ raises outliers and yields smaller clusters while increasing θ reduces outliers and yields larger clusters.

### Measuring purity of clustered FunFams

To estimate whether the clustering of an impure FunFam led to more consistent sub-families, we calculated the percentage of pure clusters. Clusters with no EC annotation were excluded. For each FunFam, we calculated the clusters with one single EC annotation as percentage of all clusters with EC annotations and defined this measure as the purity of a FunFam (Eqn. 4). We then defined the percentage of completely pure FunFams as the percentage of FunFams with a purity of 100.

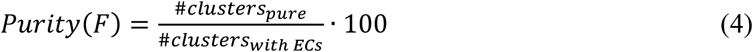

We also calculated the purity of a FunFam in terms of its size, i.e., the number of sequences contained in it:

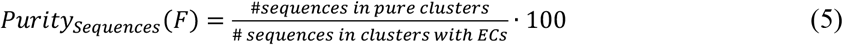

### Confidence intervals (CIs)

95% symmetric confidence intervals (CIs) were calculated from 1, 000 bootstrap samples to indicate the spread of data and certainty of average values.

### Final dataset

To construct the dataset used in this analysis, we extracted all superfamilies with at least one impure FunFam, i.e., at least one FunFam with more than one EC annotation. Since embeddings could only be computed for continuous sequences, we excluded sequences with multiple segments. After this removal, some FunFams became orphans (consisting of only a single sequence) and were also excluded. This led to a final dataset of 458 superfamilies (10.6% of all superfamilies) with 110,876 FunFams (52.1%) and 13,011 (6.1%) with EC annotations. Those 13,011 FunFams accounted for 20% of all proteins in the FunFams (1,669,245 sequences). All FunFams in a superfamily were used to determine the clustering distance cut-off θ. however, only FunFams with EC annotations were clustered to save computer time, and hence, energy.

### Availability

The final dataset and the corresponding embeddings as well as the source code used for clustering are publicly available via GitHub: https://github.com/Rostlab/FunFamsClustering. In addition, ProtBERT and PB-Tucker embeddings for any sequence can be retrieved using the bio_embeddings pipeline [42].

## Results & Discussion

### Embedding clusters increased EC purity

We began with 13,011 FunFams (6% of all) with at least one EC annotation. Of these, 1,273 (10%) contained more than one EC annotation (impure FunFams). Applying DBSCAN to all EC annotated FunFams, we split the 13,011 into 26,464 clusters (21,546 for pure and 4,918 for impure FunFams). On average, 4.4%-4.6% (95% confidence interval, CI) of the proteins in a FunFam were classified as outliers (Table S4 in Supplementary Online Material (SOM)). 63% of the DBSCAN clusters contained proteins with EC annotations; only 4% of those contained multiple EC annotations (vs. 10% in all FunFams; Fig. 1A). Only 10% of all proteins (155,044 of 1,593,567) belonged to clusters with multiple EC annotations compared to 21% (356,565 of 1,668,273) for FunFams (Fig. 1B). Thus, a larger fraction of clusters was pure (i.e., contained one EC annotation) than of FunFams both in terms of numbers of clusters and numbers of proteins (Fig. 1).

**Fig. 1:**
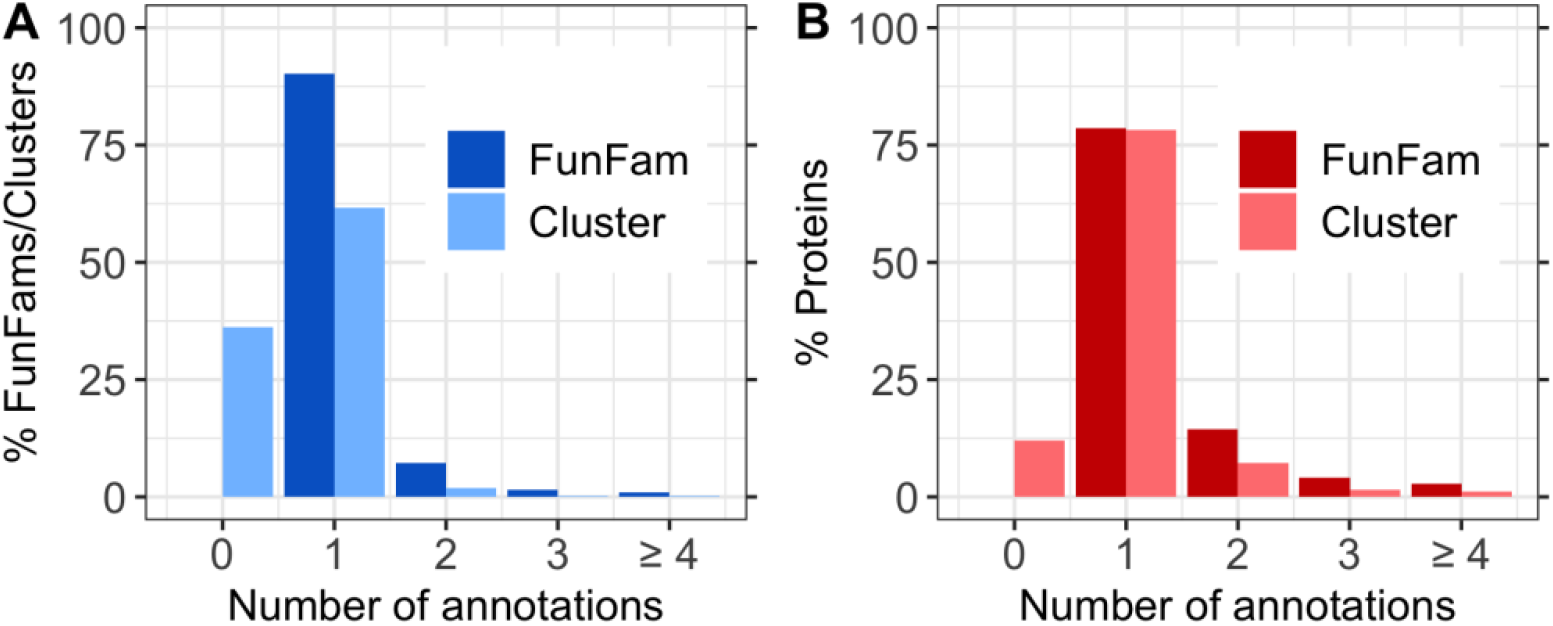
EC purity for FunFams and embedding clusters. This analysis considered 13,011 FunFams with EC annotations. Panel **A** shows the distribution of all families (FunFam/Clusters), i.e., the percentage of FunFams and embedding-based clusters with *n* EC annotations (n≥1 for FunFams and n?0 for new clusters; note: bars left and right of integer values *n*, not separated by a white space denote *n* annotations). Panel **B** shows the distribution of all proteins, i.e., the percentage of proteins in families (FunFam/Cluster) with *n* EC annotations. This number does not reveal how many proteins have an EC annotation. Of the 13,011 FunFams, 10% were impure, i.e., had multiple EC annotations (100-value for dark blue bar at 1 in panel A), and 21% of all proteins were part of these impure FunFams. After embedding-based split of FunFams, 64% (16,906) of the resulting clusters contain ECs (100-light blue bar at 0) and 4% (606) of those 16,906 were annotated to more than one EC accounting for 11% of proteins in clusters with ECs.

To further understand the extent to which the clustered FunFams provide a functionally more consistent subset, we determined for each impure FunFam, the fraction of clusters that were pure (Methods). To begin: 37% of all clusters had no EC annotated proteins and were excluded from further analysis. Of the remaining 16,906 clusters (63%), 22% were impure, i.e., contained more than one EC annotation. On average, 63% (CI: [60%; 66%]) of the clusters for a FunFam were pure (Fig. 2; dashed blue line) accounting for 58% (CI: [55%; 61%]) of all proteins (Fig. 2; dotted red line). 52% of all impure FunFams were split into completely pure clusters, i.e., for every other FunFam, the embedding-split clustered into functionally consistent sub-families (Fig. 2, right most blue point “100% Pure Clusters”) accounting for 38% of all proteins (Fig. 2, right most red point). This measure gave conservative estimates as it only considered completely pure clusters, ignoring improvements through reduction of EC annotations, e.g., when a group had originally *m+1* annotations and the clustering improved to *m*, this improvement was ignored for all *m>1*.

**Fig. 2:**
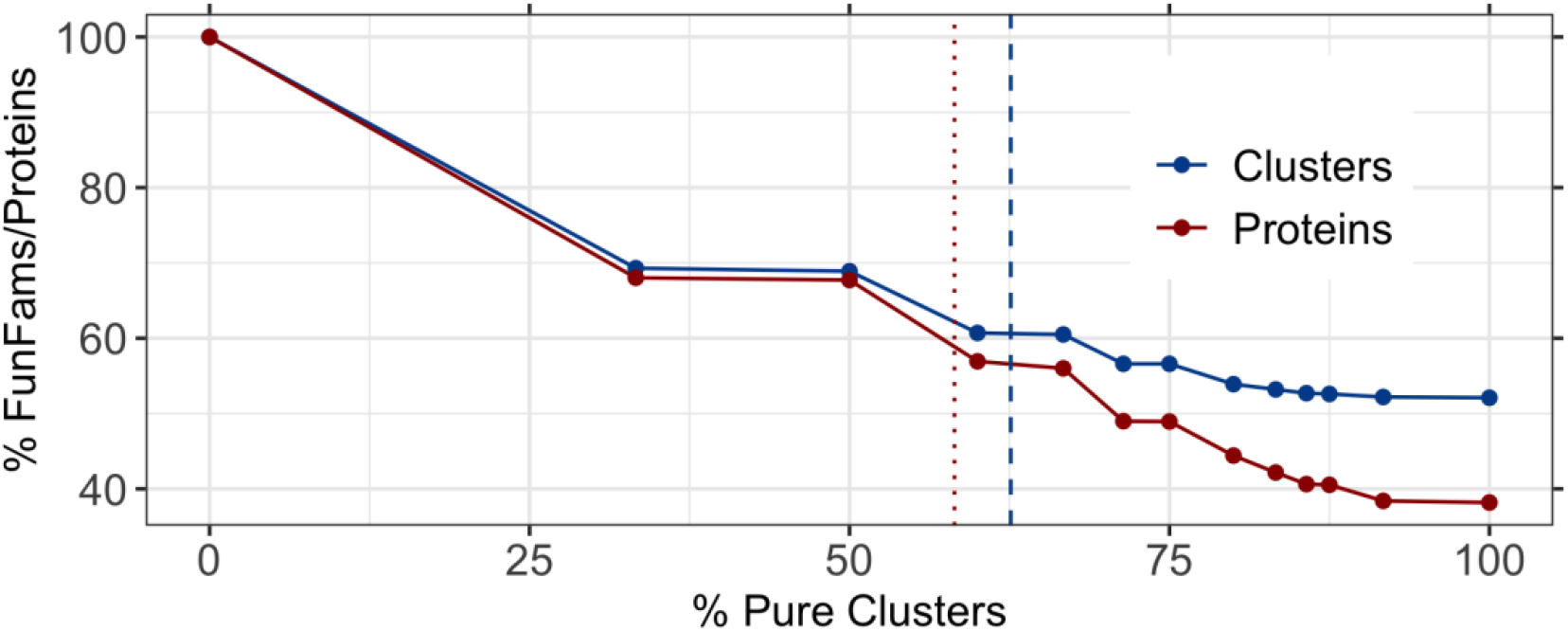
Fraction of pure clusters for impure FunFams. The y-axis gives the percentages of all clusters (blue line) or of all proteins (red line) in FunFams at levels of increasing cluster purity (Eqns. 4, 5). On average, 63% of the clusters for a FunFam were pure (dashed vertical blue line) accounting for 58% of the proteins (dotted vertical red line). 52% of the impure FunFams were split only into pure clusters (right most blue point) accounting for 38% of the proteins

### Improving EC purity without over-splitting

While splitting impure FunFams through embedding-based clustering clearly improved the EC purity, we wanted to avoid over-splitting. Trivially, the more and smaller clusters, the more likely they are pure. In the non-sense extreme of N clusters for N sequences, all clusters are trivially pure. One constraint to avoid generating too many clusters (over-splitting) is to do substantially better than by randomly splitting into the same number of clusters. We computed the random clustering using the same cluster sizes and outlier numbers as realized by the embedding-based clustering. Fewer than half as many embedding-based clusters were impure than for random (Fig. 3A, 47±1 % vs 22±1 %). Similarly, the average purity of a FunFam was almost double for embedding than for random clustering (Fig. 3B, 63±3% vs. 38±5%), and 3.5 times more FunFams were split into exclusively pure clusters by the embedding-based clustering (Fig. 3C, 52±3% vs 15±1%). This corresponded to 4.8% (CI: [4.4%; 5.2%]) of all proteins clustered into pure clusters at random compared to 38% (CI: [33%; 43%]) of all proteins for embedding-based clustering, i.e., an over 7-fold increase (Fig. 3C, red bars).

**Fig. 3:**
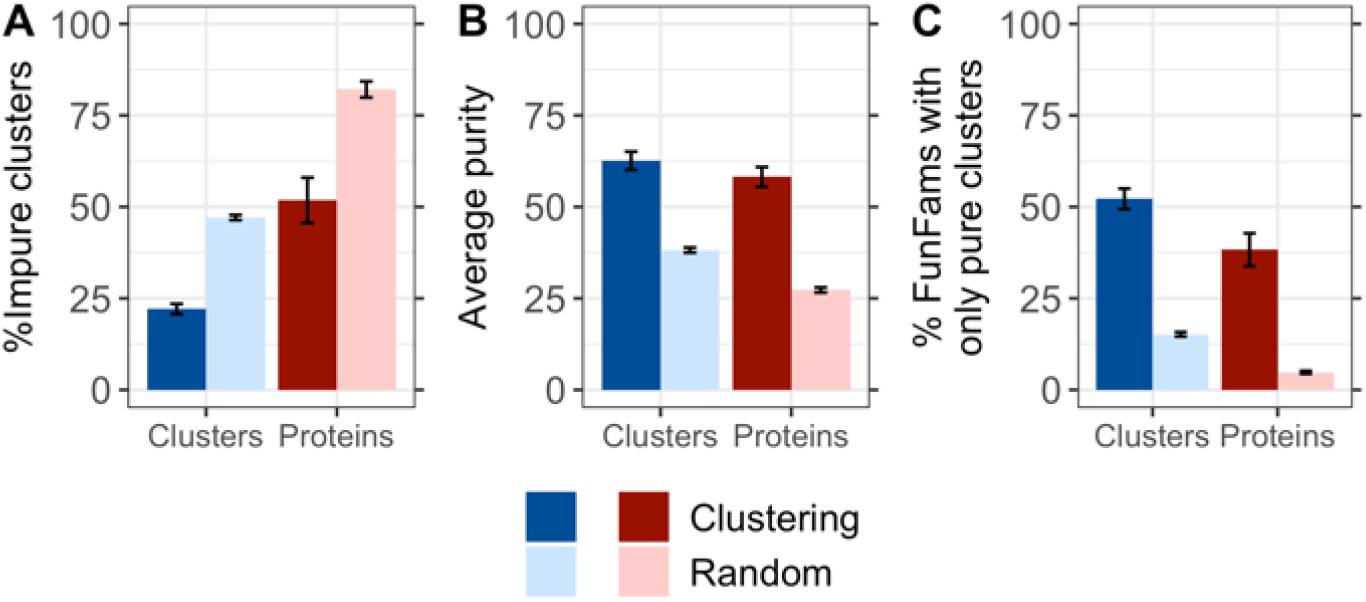
Embedding-based clusters improve EC purity over random. Random clusters were computed using the same cluster size and outlier number realized by the embedding-based clustering, but the FunFam members were randomly assigned. **A.** The fraction of impure clusters was higher for the random clustering than for our clustering (29% vs 12%). **B.** Through DBSCAN embedding-based clustering, each impure FunFam was, on average, split into 63% pure clusters while for the random clustering, the average purity was only 38%. **C.** More than half of all FunFams (53%) were split only into pure clusters for embedding-based clustering but only 15% for a random clustering. Error bars indicate symmetric 95% confidence intervals. Blue colored bars indicate numbers in terms of clusters, red colored bars in terms of proteins in those clusters. Darker colors indicate values for the clustering while lighter colors indicate values for the random approach.

An ideal split of impure FunFams generates clusters defined by two features: all members share the same EC annotation(s) and all proteins with the same EC annotation(s) are in the same cluster. Ignoring the latter leads to over-splitting. For the embedding-based clustering, 81% of the ECs occurred in one cluster (Fig. 4). However, some of the outliers had EC annotations. When also counting those (as single member clusters), the percentage of EC-exclusive clusters dropped to 63% (Fig. 4). Thus, the embedding-based clustering largely avoided over-splitting. Nevertheless, 8% of all experimentally known EC numbers were annotated to proteins from at least three different clusters (17% if including outliers; Fig. 4) and some (10%) of the outliers shared the EC number with the cluster from which they had been removed. This might indicate over-splitting or suggest a more fine-grained functional distinction between those proteins than is captured in the fourth EC level.

**Fig. 4:**
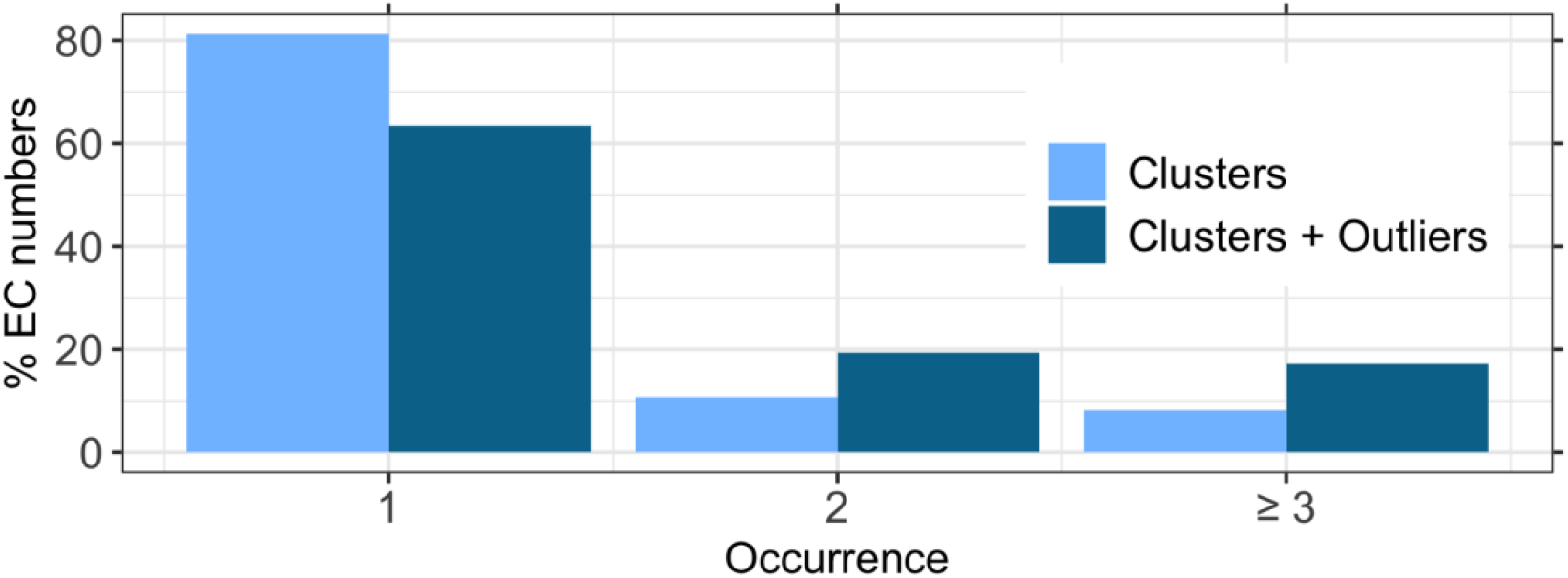
Most EC numbers only occur in one cluster. For each EC number in a FunFam, we counted the number of embedding-based clusters in which it occurred to gauge potential over-splitting. 81% of the ECs only occurred in one cluster (darker bars). If we considered outliers as clusters with one member, this number dropped to 63%. These results suggest that the clustering did not over-split the FunFams and that functionally related proteins ended up in the same cluster.

If the increased purity through clustering had been a random effect, the embedding space clustering would be EC-independent. If so, we expect no difference in the distributions of embedding distances between pure and impure FunFams, and a similar number of clusters and outliers. However, pure FunFams were, on average, split into only two clusters, while impure FunFams were split into four clusters (Table S4). This finding suggested that if a FunFam is split into many clusters, it should be considered for further manual inspection to establish whether all proteins were correctly assigned to this functional family (SOM_2.1).

### Different levels of EC annotations gave similar results

Up to this point, we only distinguished whether two proteins had the same EC annotation or not, ignoring that two proteins with ECs A.B.C.X and A.B.C.Y more likely have similar molecular function than a pair with A.* and D.*. Pairs of the first type (difference only in 4th level) will, on average, be more sequence similar than pairs of the second type (difference in 1st level). Most impure FunFams were impure due to differences on the fourth level of EC annotations (Fig. S5). Although we analyzed the clustering at higher levels of the EC classification, the results were inconclusive, probably due to data sparsity (SOM_2.2). The results for five specifically chosen superfamilies were similar (Fig. S7) underlining the more general findings that the level of EC annotation causing impurity did not crucially affect the embedding-based clustering (SOM_3.2). Instead, the performance was likely impacted more by other factors such as the presence of multifunctional enzymes and by missing annotations, i.e., an insufficient coverage by proteins with explicit experimental evidence. We applied a rather conservative definition of purity: If one protein were annotated to two EC numbers (EC1+EC2) and another protein in the same cluster were only annotated to one of those two (EC1), we would consider this cluster impure. If this difference were not due to missing annotations, we could group the two into two separate families (those with EC1+EC2, those with only EC1).

Although considering those FunFams or clusters as impure is too strict, not doing this is too loose because it will raise the performance of the random approach given that many families have many EC numbers. More importantly, the EC1-only proteins may have annotations that are incomplete or incorrect.

### Slightly worse results for experimentally verified annotations

Not all EC annotations in UniProt are experimentally verified. Only using experimentally verified EC annotations (with evidence code ECO:0000269) for sequences in Swiss-Prot, 4,709 FunFams contained any sequence with EC annotations and 637 (14%) were impure.

46% of those impure FunFams were clustered into fully pure clusters, and our approach achieved an average purity of 53% being slightly worse than for the full set. This could be due to missing annotations: If two proteins A and B in a family or cluster are annotated to the same two EC numbers EC1 and EC2, but for protein A only EC1 is experimentally verified and for protein B EC2, such a family or cluster would be considered impure. This explanation is also supported by the fact that a larger fraction of FunFams was impure if only experimentally verified annotations were considered (14% vs 10%).

### Similar improvement for single domain proteins

Most EC annotations are available for an entire protein, not for the structural domain responsible for this function, yet most proteins have several domains [43]. To avoid the potential problem of multi-domain proteins for purity, we assessed the clustering results for 2,412 FunFams only consisting of single-domain proteins. Of those, 6% (136) were impure vs. 10% for all FunFams. This provided a rough idea for the problem of our purity definition: roughly half (10/6~2) of the problem related to multi-domain proteins.

Clustering single domains performed similarly to full-length proteins: 52% of the impure FunFams with single-domain proteins were clustered into fully pure clusters vs. 52% for the entire set, and the clustering achieved an average purity of 60% (vs. 63%). These results showed that the clustering approach did not only remove the impurity caused by inaccurate annotations but could identify functional relationships not detected using sequence comparison.

### Details of parameter choice mattered

For a more detailed analysis of particular details of our method, in particular, for the choice of embeddings and clustering parameters, we chose five superfamilies with diverse properties (CATH identifiers: 3.40.50.150, 3.20.20.70, 3.40.47.10, 3.50.50.60, 1.10.630.10; SOM_3).

#### Embedding space better captured functional sub-groups than sequence space

Instead of using embedding distance for DBSCAN, we redid the clustering using the pairwise percentage sequence identity (PIDE) between any two sequences in a FunFam, converted into a sequence distance (1-PIDE). This approach generated more clusters with more outliers that were, on average, less pure than from embeddings (Table S7). This implied that embedding similarity captured functional relationships between sequences better than sequence similarity.

#### Euclidean distance yielding best clustering results

Cosine distance is often considered more standard for comparing embeddings, e.g., in NLP. Replacing Euclidean distance with Cosine distance (Eqn. 2) revealed that most embeddings of proteins within a FunFam were represented by vectors with the same orientation leading to a Cosine distance of 0, i.e., we could not use it to split any FunFam (Table S7). Euclidean distance often suffers more from problems with high dimensional spaces [44] than Manhattan distance (Eqn. 3). Indeed, for our problem Manhattan, unlike cosine, distance worked but not better than Euclidean (Table S7). Since PB-Tucker was optimized on Euclidean distance, we clustered on Euclidean.

#### PB-Tucker embeddings largely superior to ProtBERT embeddings

In direct comparison between ProtBERT and PB-Tucker embeddings, we observed the following when clustering the five superfamilies (Table S3): the number of clusters and outliers was slightly smaller for ProtBERT, but the fraction of impure clusters was higher than for PB-Tucker (19% for ProtBERT vs 13% for “default”; Table S7). The average purity was also higher for PB-Tucker (“default” = 59%) than for ProtBERT (51%; Table S7). Thus, PB-Tucker appeared superior in capturing functional differences.

#### Smaller distance thresholds resulted in smaller and purer clusters

The distance threshold θ of DBSCAN defines whether or not two points are close enough to each other to be grouped. For the default clustering, we chose the median distance between all proteins for each superfamily (Methods). The observed distribution of distances (Fig. S3) suggested choosing superfamily-specific thresholds. As expected, the smaller θ, the more clusters and outliers will result (“θ =1^st^ quartile” vs “default”; Table S7). Largely due to splitting FunFams into more clusters at smaller θ, the resulting clusters were seemingly purer with only 4% impure clusters (vs. 13%) and an average purity of 83% (vs. 59%; Table S7). In contrast, larger θ thresholds (here the 3^rd^ quartile) affected fewer, more impure clusters (Table S7). Thus, the choice of θ highly influenced the clustering results. For some applications, lower values of θ might be best to obtain a large, highly consistent set of small sub-families that can serve e.g., as seed to further extend those sub-families to larger functionally related families. Also, especially for FunFams for which using a larger cutoff largely failed, decreasing the distance threshold can help to still identify which sequences might cause impurity.

#### Default neighborhood size resulted in best clustering

DBSCAN forms clusters around “core points” which are points with at least *n* neighbors. For the five superfamilies, we tested fixed neighborhood sizes of *n*∈[5;129;255] and variable neighborhood sizes dependent on the size of the FunFam *n*=x*|F|, with |F| as the number of proteins in a FunFam and x∈[0.01;0.1;0.2] (Methods). While *n*=129 and *n*=255 were in the range of what is recommended for *n* [41], the clustering was worse than for the default parameter (*n*=5) (Table S7). Specifically, the number of outliers exploded for these large neighborhood sizes (Table S7, Fig. S6).

Since FunFams differ substantially in the number of proteins, we hypothesized that – similar as for θ – it could be reasonable to choose a different *n* for each FunFam. However, this did not improve compared to the default clustering (Table S7); the default *n*=5 was a good choice.

### Clustering increased purity of ligand binding

Another way to assess the purity of molecular protein function in protein families is to compare their similarity in terms of ligand-binding. We extracted bound ligands from BioLip [45] and only considered annotations defined as the cognate ligand [46] (SOM_1.3). Of the 13,011 FunFams considered so far, 950 (7%) contained any annotation about a ligand bound, and of those 950, 158 (17%) were annotated with more than one unique ligand. Embedding-based clustering split 33% of these FunFams into clusters with only one type of ligand, i.e., “pure” clusters (compared to 52% for EC level 4) and an average purity of 36% (compared to 63% for ECs). Although ligand annotations remained limited, these results confirmed that embedding-based clustering increased functional purity of FunFams for an aspect of function not used during method development.

### Increased purity for non-FunFam sequence families

We expected the approach to succeed for any protein family grouping. To establish this, we clustered five superfamilies (Table S4) at 35% PIDE. This resulted in 2,160 families – dubbed S35 – not optimized for function (SOM_1.4); 116 of the S35 were impure. Embedding-based clustering resulted in 35±9% fully pure clusters and an average purity of 46±9% (Fig. 5B&C, darker blue bars). Clustering by sequence distance (1-PIDE) led to only 31±9% fully pure clusters and an average purity of 42±9% (Fig. 5B&C, lighter blue bars). While the difference between sequence- and embedding-based clustering was less than for FunFams, the comparison was very conservative in that the construction of the S35 implicitly favored sequence similarity. Splitting the S35 families into consistent functional clusters worked surprisingly well compared to FunFams because the S35 were less well grouped by function than FunFams, i.e., some functional inconsistency was easy to identify. Given that the construction of the S35 made no other assumption than relation by sequence (PIDE), this experiment provided a rather conservative proof-of-principle that embedding-based clustering is likely to outperform sequence-based clustering for any grouping of proteins.

**Fig. 5:**
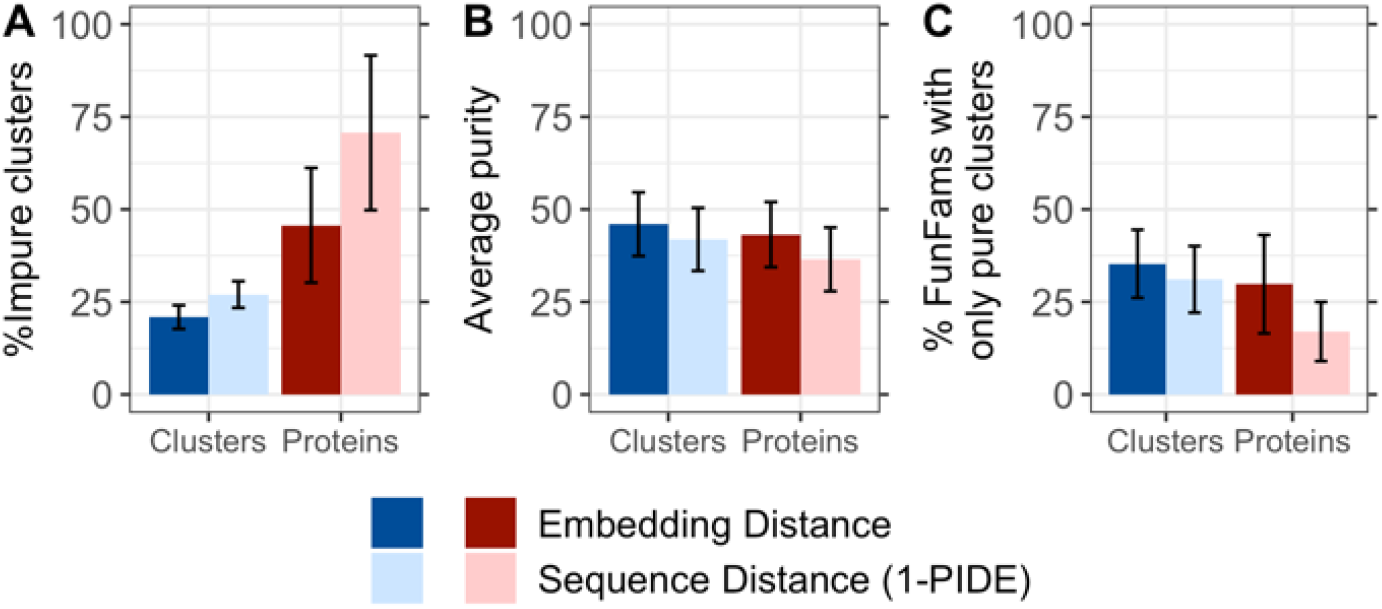
Clustering results for S35 families. S35 families were created by clustering CATH superfamilies at 35% sequence identity resulting in 116 impure S35 families. Those were clustered using sequence distance (1-PIDE) and embedding distance. **A.** The fraction of impure clusters was higher for the clustering using sequence distance than for the clustering based on embeddings (27% vs 21%). **B.** Through embedding-based clustering, each impure S35 family was, on average, split into 46% pure clusters while for the sequence-based clustering, the average purity was 42%. **C.** 35% of impure S35 families were split only into pure clusters for embedding-based clustering and 31% for sequence-based clustering. Error bars indicate symmetric 95% confidence intervals. Blue colored bars indicate numbers in terms of clusters, red colored bars in terms of proteins in those clusters. Darker colors indicate values for the clustering while lighter colors indicate values for the random approach.

## Conclusions

FunFams (Das, et al., 2016; Sillitoe, et al., 2013) provide a high-quality sub-classification of CATH superfamilies into families of functionally related proteins (Sillitoe, et al., 2020; Sillitoe, et al., 2019). However, some FunFams are impure and 7% of all FunFams with EC annotations contain at least two different ECs (Fig. S1). Here, we introduced a novel approach toward clustering proteins through embeddings derived from the LM ProtBERT (Elnaggar, et al., 2020) and further optimized to capture relationships between proteins within one CATH superfamily (called *PB-Tucker*). Similarity between embeddings can capture information different from what is captured by sequence similarity. In particular, it can reveal new functional relations between proteins (Asgari and Mofrad, 2015; Hamid and Friedberg, 2019; Littmann, et al., 2021; Vig, et al., 2020; Villegas-Morcillo, et al., 2020; Yang, et al., 2018). Clustering all FunFams with more than one EC annotation (impure FunFams) using DBSCAN (Ester, et al., 1996) reduced the percentage of impure clusters to 22% (CI: [21%, 23%]). An impure FunFam was on average split into 63% pure clusters (CI: [60%: 66%]) and more than half (53%, CI: [50%; 56%]) of all impure FunFams were split into fully pure sub-families (Fig. 2). This corresponded to a four-fold increase over random clustering (Fig. 3B). In terms of number of proteins (rather than number of clusters), the increase was almost ten-fold. Only 4.8% (CI: [4.4%; 5.2%]) of the proteins were in FunFams split into pure clusters for random while this number rose to 38% (CI: [33%; 43%]) for the *PB-Tucker* embedding-based clustering.

A more detailed analysis of five hand-picked superfamilies (Table S5) showed that the default choices for the DBSCAN parameters were reasonable (Fig. S6, Table S7; with the default *n*=5 and the distance threshold θ determined automatically. Also, Euclidean distance between embeddings worked best (Table S7).

Restricting the analysis to manually curated EC annotations limited the validation of our approach to a small fraction (6.1%) of all FunFams and even for those FunFams, most EC annotations remain unknown. Nevertheless, we have shown that our approach could capture more fine-grained functional relationships and enabled splitting FunFams into more functionally consistent sub-families. Especially for FunFams without many known functional annotations, our clustering can be used to (i) investigate whether or not the family could be impure based on the number of clusters resulting from the embedding-based split, or (ii) more safely infer functional annotations between members of one functional cluster than between members of one FunFam. We presented evidence suggesting that the findings for EC annotations will hold for other aspects of protein function, e.g., for binding. While we only applied this approach to FunFams using embeddings optimized for CATH, this clustering could be applied to any database of functional families using a more generalized version of those embeddings.

## Abbreviations

DBSCAN: density-based spatial clustering of applications with noise
d: dimensions
EC: Enzyme Commission
FunFam: functional family
LM: language model
NLP: natural language processing
PIDE: percentage sequence identity

## Acknowledgements

Thanks to Tim Karl and Inga Weise (both TUM) for invaluable help with technical and administrative aspects of this work. We would like to acknowledge Ian Sillitoe (UCL) for helpful comments on EC data. Last, but not least, thanks to all maintainers of public databases and to all experimentalists who enabled this analysis by making their data publicly available.

This work was supported by the Bavarian Ministry of Education through funding to the TUM, by a grant from the Alexander von Humboldt foundation through the German Ministry for Research and Education (BMBF: Bundesministerium für Bildung und Forschung), and by two grants from BMBF (031L0168 and Program “Software Campus 2.0 (TUM): 01IS17049) as well as by a grant from Deutsche Forschungsgemeinschaft (DFG–GZ: RO1320/4–1). We gratefully acknowledge the support of NVIDIA Corporation with the donation of one Titan GPU used for this research. Nicola Bordin acknowledges financial support from the Biotechnology and Biological Sciences Research Council (UK) [BB/R009597/1].

## References

[1] S. S. Hannenhalli and R. B. Russell, “Analysis and prediction of functional sub-types from protein sequence alignments,” J Mol Biol, vol. 303, no. 1, pp. 61–76, Oct. 2000, doi: 10.1006/jmbi.2000.4036.

[2] C. A. Orengo, A. D. Michie, S. Jones, D. T. Jones, M. B. Swindells, and J. M. Thornton, “CATH--a hierarchic classification of protein domain structures,” Structure, vol. 5, no. 8, pp. 1093–108, Aug. 1997, doi: 10.1016/s0969-2126(97)00260-8.

[3] “CATH: Protein Structure Classification Database at UCL.”https://www.cathdb.info/ (accessed Nov. 02, 2020).

[4] I. Sillitoe et al., “CATH: increased structural coverage of functional space,” Nucleic Acids Res., no. gkaa1079, Nov. 2020, doi: 10.1093/nar/gkaa1079.

[5] S. Das, D. Lee, I. Sillitoe, N. L. Dawson, J. G. Lees, and C. A. Orengo, “Functional classification of CATH superfamilies: a domain-based approach for protein function annotation,” Bioinformatics, vol. 32, no. 18, p. 2889, Sep. 2016, doi: 10.1093/bioinformatics/btw473.

[6] I. Sillitoe et al., “New functional families (FunFams) in CATH to improve the mapping of conserved functional sites to 3D structures,” Nucleic Acids Res, vol. 41, no. Database issue, pp. D490–8, Jan. 2013, doi: 10.1093/nar/gks1211.

[7] N. Zhou et al., “The CAFA challenge reports improved protein function prediction and new functional annotations for hundreds of genes through experimental screens,” Genome Biol, vol. 20, no. 1, p. 244, Nov. 2019, doi: 10.1186/s13059-019-1835-8.

[8] S. Das et al., “CATH FunFHMMer web server: protein functional annotations using functional family assignments,” Nucleic Acids Res, vol. 43, no. W1, pp. W148–53, Jul. 2015, doi: 10.1093/nar/gkv488.

[9] S. Das, H. M. Scholes, and C. A. Orengo, “CATH functional families predict protein functional sites,” 2020, doi: 10.1101/2020.03.23.003012.

[10] L. Scheibenreif, M. Littmann, C. Orengo, and B. Rost, “FunFam protein families improve residue level molecular function prediction,” BMC Bioinformatics, vol. 20, no. 1, p. 400, Jul. 2019, doi: 10.1186/s12859-019-2988-x.

[11] E. C. Webb, Enzyme nomenclature 1992: recommendations of the Nomenclature Committee of the International Union of Biochemistry and Molecular Biology on the nomenclature and classification of enzymes. Academic Press, 1992.

[12] C. J. Jeffery, “Moonlighting proteins: old proteins learning new tricks,” Trends Genet, vol. 19, no. 8, pp. 415–7, Aug. 2003, doi: 10.1016/S0168-9525(03)00167-7.

[13] M. Heinzinger et al., “Modeling aspects of the language of life through transfer-learning protein sequences,” BMC Bioinformatics, vol. 20, no. 1, p. 723, Dec. 2019, doi: 10.1186/s12859-019-3220-8.

[14] R. Rao et al., “Evaluating Protein Transfer Learning with TAPE,” ArXiv190608230 Cs Q-Bio Stat, Jun. 2019, Accessed: Nov. 03, 2020. [Online]. Available: http://arxiv.org/abs/1906.08230.

[15] E. C. Alley, G. Khimulya, S. Biswas, M. AlQuraishi, and G. M. Church, “Unified rational protein engineering with sequence-based deep representation learning,” Nat. Methods, vol. 16, no. 12, Art. no. 12, Dec. 2019, doi: 10.1038/s41592-019-0598-1.

[16] A. Elnaggar et al., “ProtTrans: Towards Cracking the Language of Life’s Code Through Self-Supervised Deep Learning and High Performance Computing,” bioRxiv, p. 2020.07.12.199554, Jul. 2020, doi: 10.1101/2020.07.12.199554.

[17] A. Madani et al., “ProGen: Language Modeling for Protein Generation,” bioRxiv, p. 2020.03.07.982272, Mar. 2020, doi: 10.1101/2020.03.07.982272.

[18] M. E. Peters et al., “Deep contextualized word representations,” ArXiv180205365 Cs, Mar. 2018, Accessed: Nov. 02, 2020. [Online]. Available: http://arxiv.org/abs/1802.05365.

[19] A. Radford, K. Narasimhan, T. Salimans, and I. Sutskever, “Improving Language Understanding by Generative Pre-Training,” p. 12.

[20] J. Devlin, M.-W. Chang, K. Lee, and K. Toutanova, “BERT: Pre-training of Deep Bidirectional Transformers for Language Understanding,” ArXiv181004805 Cs, May 2019, Accessed: Nov. 02, 2020. [Online]. Available: http://arxiv.org/abs/1810.048051.

[21] A. Rives et al., “Biological structure and function emerge from scaling unsupervised learning to 250 million protein sequences,” bioRxiv, p. 622803, Jan. 2020, doi: 10.1101/622803.

[22] M. Littmann, M. Heinzinger, C. Dallago, T. Olenyi, and B. Rost, “Embeddings from deep learning transfer GO annotations beyond homology,” bioRxiv, p. 2020.09.04.282814, Oct. 2020, doi: 10.1101/2020.09.04.282814.

[23] M. Ester, H.-P. Kriegel, J. Sander, and X. Xu, “A density-based algorithm for discovering clusters in large spatial databases with noise,” 1996, vol. 96, pp. 226–231.

[24] D. A. Lee, R. Rentzsch, and C. Orengo, “GeMMA: functional subfamily classification within superfamilies of predicted protein structural domains,” Nucleic Acids Res., vol. 38, no. 3, pp. 720–737, Jan. 2010, doi: 10.1093/nar/gkp1049.

[25] M. Steinegger, M. Meier, M. Mirdita, H. Vöhringer, S. J. Haunsberger, and J. Söding, “HH-suite3 for fast remote homology detection and deep protein annotation,”BMC Bioinformatics, vol. 20, no. 1, p. 473, Sep. 2019, doi: 10.1186/s12859-019-3019-7.

[26] J. A. Capra and M. Singh, “Characterization and prediction of residues determining protein functional specificity,”Bioinformatics, vol. 24, no. 13, pp. 1473–1480, Jul. 2008, doi: 10.1093/bioinformatics/btn214.

[27] T. U. Consortium, “UniProt: a worldwide hub of protein knowledge,”Nucleic Acids Res., vol. 47, no. D1, pp. D506–D515, Jan. 2019, doi: 10.1093/nar/gky1049.

[28] “UniProt.”https://sparql.uniprot.org/ (accessed Nov. 12, 2020).

[29] D. Bahdanau, K. Cho, and Y. Bengio, “Neural Machine Translation by Jointly Learning to Align and Translate,”ArXiv14090473 Cs Stat, May 2016, Accessed: Nov. 02, 2020. [Online]. Available: http://arxiv.org/abs/1409.0473.

[30] D. Shen et al.,“Baseline Needs More Love: On Simple Word-Embedding-Based Models and Associated Pooling Mechanisms,”ArXiv180509843 Cs, May 2018, Accessed: Nov. 02, 2020. [Online]. Available: http://arxiv.org/abs/1805.09843.

[31] F. Pedregosa et al.,“Scikit-learn: Machine Learning in Python,”J. Mach. Learn. Res., vol. 12, pp. 2825–2830, 2011.

[32] J. Sander, M. Ester, H.-P. Kriegel, and X. Xu, “Density-Based Clustering in Spatial Databases: The Algorithm GDBSCAN and Its Applications,”Data Min. Knowl. Discov., vol. 2, no. 2, pp. 169–194, Jun. 1998, doi: 10.1023/A:1009745219419.

[33] J. Yang, A. Roy, and Y. Zhang, “BioLiP: a semi-manually curated database for biologically relevant ligand–protein interactions,”Nucleic Acids Res., vol. 41, no. Database issue, pp. D1096–D1103, Jan. 2013, doi: 10.1093/nar/gks966.

[34] J. D. Tyzack, L. Fernando, A. J. M. Ribeiro, N. Borkakoti, and J. M. Thornton, “Ranking Enzyme Structures in the PDB by Bound Ligand Similarity to Biological Substrates,”Structure, vol. 26, no. 4, pp. 565–571.e3, Apr. 2018, doi: 10.1016/j.str.2018.02.009.

