## Supporting Online Material for "Clustering FunFams using sequence embeddings improves EC purity"

#### Table of Contents for Supporting Online Material

### Short description of Supporting Online Material

In this Supporting Online Material (SOM), we provide more details on the underlying language models used to create protein embeddings and on how binding annotations were obtained. Also, we show a more detailed analysis of the results discussed in the main text.

Of the 22,830 FunFams with EC annotations, 7% (1,526) are impure accounting for 16% of all sequences (Fig. S1). Of the 92,024 proteins with EC annotations, 8% (7,586) are multifunctional, i.e., annotated to more than one EC number (Table S1). PB-Tucker embeddings were derived from ProtBERT embeddings by optimizing them for distinguish CATH classes (Fig. S2: Visualization of the embedding generation using ProtBERT and PB-TuckerFig. S2). Comparing the performance of ProtBERT and PB-Tucker embeddings for the inference of CATH class and comparing their distance distribution for five superfamilies showed that PB-Tucker seemed to be the better choice for our approach (Table S2, Fig. S3).

To assess whether our clustering approach also worked for protein classifications other than FunFams, we clustered five superfamilies at 35% sequence identity and dubbed the resulting alignment S35 families (Table S3).

Using PB-Tucker and DBSCAN to cluster 13,011 FunFams with EC annotations, we observed different results for pure and impure FunFams (Table S4, Fig. S4). Assessing the clustering performance for different EC levels did not show a clear trend (Fig. S5) indicating that the approach worked for functionally very different and more similar sequences. To further investigate the influence of certain parameters, we chose five superfamilies for five different criteria (Table S5). Testing different distance thresholds and neighborhood sizes for DBSCAN as well as different distance metrics could not improve over the original choice of parameters (Table S6, Table S7, Fig. S6). Assessing the clustering on different levels of EC annotations for those five superfamilies showed that our approach was more influenced by the presence of multifunctional proteins or potentially missing annotations than by the level on which ECs were different (Fig. S7).

### 1. Materials & Methods

#### 1.1. Dataset

##### Fig. S1: Fraction of FunFams with x EC annotations.

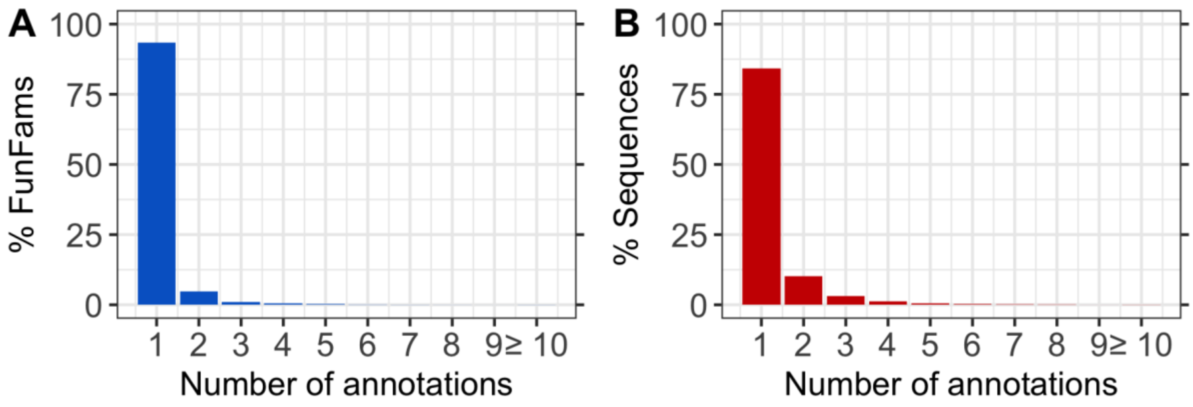

Since FunFams and EC annotations both provide a classification of proteins into functionally related classes, we expected FunFams to always only have one EC annotation. Surprisingly, some FunFams have multiple EC annotations. Panel **A** shows the fraction of FunFams with *n* EC annotations. 11% of FunFams have any EC annotation. Of those, 7% have multiple annotations. Panel **B** shows the relative size of FunFams with *n* EC annotations. The relative size is measured as the fraction of sequences that are in FunFams with *n* EC annotations. This number does not give any information about how many sequences have an EC annotation.

##### Table S1: Number of multifunctional proteins in final dataset. *

| | |
|---|---|
| **No. of proteins with EC annotations** | 92,024 |
| **No. of proteins with >1 EC number** | 7,586 (8%) |
| **No. of proteins with >1 EC number (different on the 1st level)** | 3,524 (47% of 7,586) |
| **No. of proteins with >1 EC number (different on the 2nd level)** | 548 (7%) |
| **No. of proteins with >1 EC number (different on the 3rd level)** | 684 (9%) |
| **No. of proteins with >1 EC number (different on the 4th level)** | 2,830 (37%) |

* In the final dataset, 8% of all proteins with EC annotations were annotated to more than one EC number. 37% of those were annotated to EC numbers only different on the 4th level (promiscuous enzymes).

#### 1.2. Protein representation.

The pre-trained and publicly available model ProtBERT-BFD (Elnaggar *et al.*, 2020) (in the following called *ProtBERT*) was used to create fixed-length vector representations (embeddings) for protein sequences (Fig. S2). For ProtBERT, a stack of 30 attention layers, each having 16 attention heads with a hidden state size of 1024 (total number of free parameters: 420M) was trained on BFD (Steinegger and Söding, 2018; Steinegger, Mirdita and Söding, 2019). In contrast to BERT which uses a second loss (next-sentence prediction) to train a special token that summarizes sequences of variable length, ProtBERT was solely trained on the masked language modeling loss, i.e., on reconstructing corrupted input tokens.

To create PB-Tucker embeddings, ProtBERT representations were projected in two steps from 1024-dimensions (d) first to 512-d and then further down to 128-d using a two-layer, fully connected neural network with tanh non-linearity between the layers (Fig. S2). The distance between two samples was defined as the Euclidean distance between those 128-d vectors and a soft margin loss was deployed to optimize distances between triplets in this new space. For optimization, the Adam optimizer with a learning rate of 0.001 and a batch size of 256 was used. For training, a non-redundant version of CATH v4.3 (*CATH: Protein Structure Classification Database at UCL*, 2020) clustered at 100% sequence identity (PIDE) (122,727 proteins) was clustered further at 95% PIDE and 95% coverage resulting in 66,980 proteins. Profiles were created by iteratively searching the representatives of this second clustering step against UniRef50 (Suzek *et al.*, 2015). The consensus sequences of the resulting profiles were further clustered at 20% PIDE and 50% coverage using transitive clusters (connecting clusters if any members between two clusters fulfil the clustering criteria on PIDE and coverage) to detect remote homologs leaving 10,100 proteins. A random subset of 100 proteins of the remaining 10,100 cluster representatives from different homologous superfamilies according to CATH (level H in the hierarchy) was used to determine early stopping. Training was performed using a subset of the 95% non-redundant set (66,980 proteins) so that any protein (i) was not part of any test set protein cluster and (ii) did not belong to the same homologous superfamily according to CATH as any protein in the test set resulting in a final set of 51,333 proteins for training. For evaluation, a lookup set was created the same way as the training set but allowing for proteins of the same homologous superfamily, resulting in a set of size 64,709. Profile creation and clustering was done using (Steinegger and Söding, 2017, 2018).

How well PB-Tucker learned to group proteins with similar CATH classes and differentiate between proteins from different CATH classes was assessed by measuring the accuracy of predicting all four levels of CATH through annotation transfer. For every query protein from the test set, the protein with the smallest Euclidean distance in the lookup set was identified and its CATH class transferred to the query protein. This resulted in an accuracy of 38% for predicting all four levels of the CATH hierarchy using PB-Tucker compared to only 8% when using ProtBERT (Table S2). This huge difference in performance clearly showed that PB-Tucker embeddings better capture the CATH hierarchy than ProtBERT embeddings.

**Table S2: Accuracy for inferring CATH classes using ProtBERT or PB-Tucker embeddings. ***

| | ProtBERT | PB-Tucker |
|---|---|---|
| Class | 58 | 67 |
| Architecture | 28 | 48 |
| Topology | 11 | 33 |
| Homologous Superfamily | 8 | 38 |

* The performance of ProtBERT and PB-Tucker is compared by measuring their accuracies when predicting all four levels of CATH by looking up the closest neighbor for each of the 100 test proteins in the lookup set. Especially for the lower levels of the hierarchy, PB-Tucker clearly outperforms ProtBERT, highlighting the benefit of using contrastive learning in combination with pre-trained language models.

**Fig. S2: Visualization of the embedding generation using ProtBERT and PB-Tucker.**

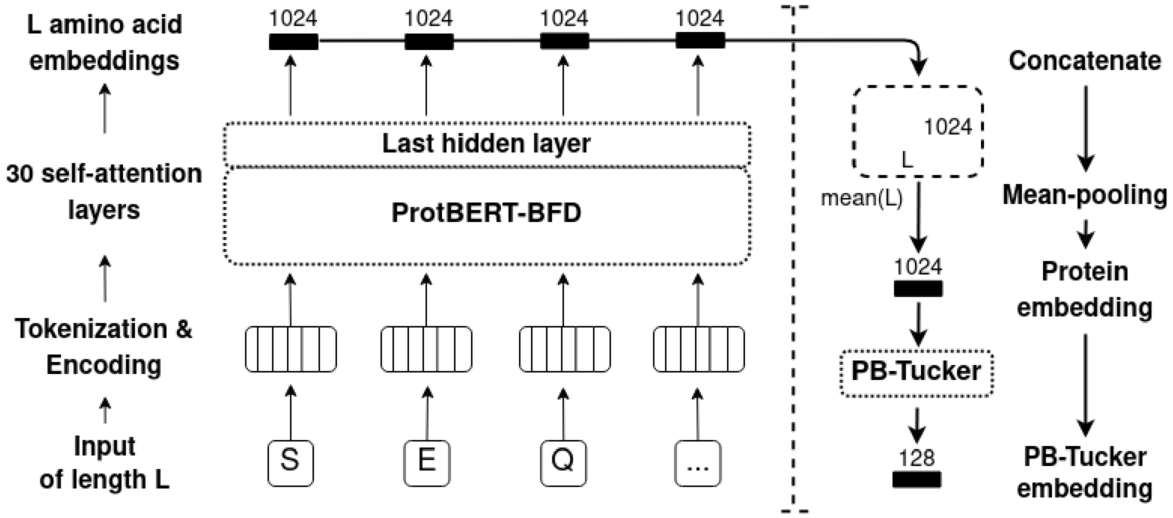

We outline the process of generating fixed-size embeddings for protein sequences of variable length using ProtBERT and PB-Tucker. The example illustrated here shows how the first three residues of protein sequence ("SEQ..") are processed: First, each residue is projected onto a vector space using an encoding layer and a stack of 30 self-attention layers. In a second step, embeddings of the last attention layer are extracted, concatenated and averaged over the length of the protein (global average pooling), resulting in a 1024-dimensional embedding. Those ProtBERT embeddings are mapped onto a new, 128-dimensional embedding using PB-Tucker which has been optimized to group embeddings of sequences similar in terms of the CATH hierarchy closer together than dissimilar proteins.

Comparing the distances between sequences within the same FunFam and those between different FunFams in the same superfamily, we observed larger relative differences for PB-Tucker (Fig. S3A) than for ProtBERT (Fig. S3B). The distribution of distances between sequences from different FunFams was narrower for most of the chosen superfamilies for PB-Tucker except for 1.10.630.10. The latter was also the superfamily with the smallest difference between within and between FunFam distances for both ProtBERT and PB-Tucker (Fig. S3). These observations suggest that PB-Tucker in fact better captures functional relationships within superfamilies and FunFams and is therefore a reasonable choice to use to further cluster FunFams.

**Fig. S3: Average sequence distances for five superfamilies.**

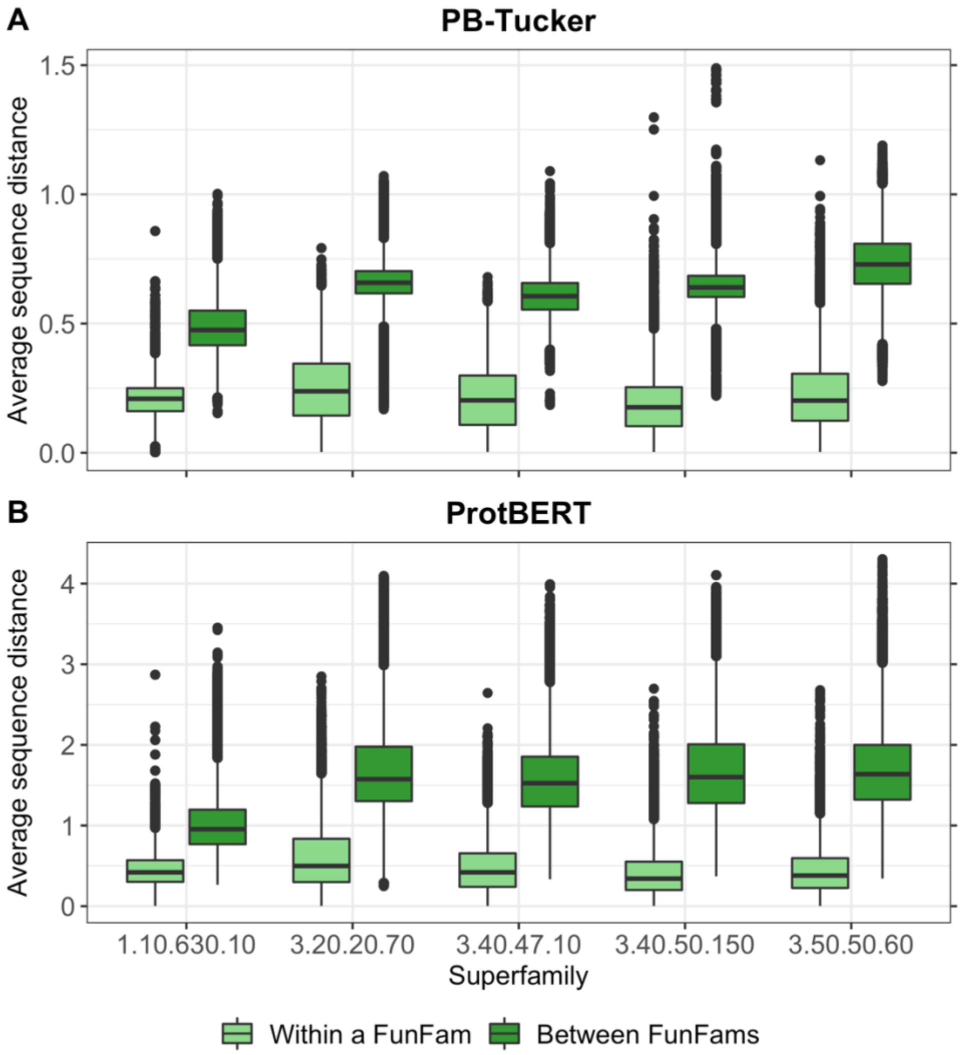

For each sequence in any FunFam of five exemplary superfamilies, we calculated the average distance of this sequence to all other sequences in this FunFam ("within a FunFam", lighter green boxes) and to sequences in other FunFams in the same superfamily ("between FunFams", darker green boxes) using **A.** PB-Tucker embeddings (128 dimensions) or **B.** ProtBERT embeddings (1024 dimensions). Grouping the resulting distances by superfamily and comparing the distributions showed that distances varied between superfamilies making it unreasonable to use the same distance cutoff for clustering for all superfamilies. The difference between within and between FunFam distances was in general larger for PB-Tucker than for ProtBERT making it reasonable to use PB-Tucker for clustering FunFams.

##### 1.3. Information on bound ligand.

We extracted annotations of bound ligands from BioLip (Yang, Roy and Zhang, 2013). BioLip provides information on ligand binding based on structural information from PDB (Berman *et al.*, 2000; *RCSB PDB*, 2020). Therefore, it is possible to have multiple annotations for one sequence if multiple structures for that sequence exist. To obtain annotations per sequence, we extracted binding information for all chains of structures matching a given sequence, which have been determined through X-ray crystallography (Smyth and Martin, 2000) with a resolution of 2.5Å and combined these annotations. It has been shown, that many ligands bound to enzymatic structures in PDB are not similar to the cognate ligand binding this enzyme under native conditions (Tyzack *et al.*, 2018). We considered only cognate ligands for our analysis. Since FunFams often only cover part of a sequence, we considered a sequence as bound to a ligand only if at least one of the corresponding binding residues was part of the sequence stretch covered in the FunFam alignment.

##### 1.4. Constructing S35 families.

We constructed S35 alignments from CATH superfamilies by clustering all sequences in one superfamily at 35% sequence identity using MMseqs2 (Steinegger and Söding, 2017, 2018). This resulted in 2,160 S35 families of which 571 contained at least one sequence with EC annotations and 116 were impure.

---

**Table S3: S35 families for five CATH superfamilies. ***

| Superfamily | Number of sequences | Number of S35 families |
|---|---|---|
| 3.40.50.150 | 34,022 | 786 |
| 3.20.20.70 | 43,900 | 365 |
| 3.40.47.10 | 7,811 | 117 |
| 3.50.50.60 | 20,022 | 546 |
| 1.10.630.10 | 9,716 | 346 |

* We clustered five superfamilies in S35 families of sequences with at least 35% sequence identity resulting in 2,160 S35 families in total.

### 2. Additional results for EC analysis of clustering

#### 2.1. Difference in clustering for pure and impure FunFams.

Pure FunFams were on average split into two clusters while impure FunFams were split into four clusters (Table S4). Especially if a FunFam is split into many clusters (e.g., ≥4; Fig. S4), this can be an indicator for functional impurity. EC annotations were not complete for most FunFams (on average only 16% of sequences in a FunFam have EC annotations). Therefore, there could exist more impure FunFams than captured through the EC purity. The number of clusters could provide a reasonable first step to identify impure FunFams that need further refinement but for which EC annotations are not available or incomplete.

---

**Table S4: Summary of clustering results for FunFams with EC annotations. ***

| | All | Pure | Impure |
|---|---|---|---|
| **Number of FunFams** | 13,011 | 11,738 (90%) | 1,273 (10%) |
| **Number of clusters (Fold increase)** | 26,464 (2.03; CI:[1.99;2.07]) | 21,546 (1.84; CI:[1.80;1.88]) | 4,918 (3.9; CI:[3.7;4.1]) |
| **Number (Fraction) of sequences classified as outliers** | 74,706 (4.5%; CI:[4.4%;4.6%]) | 59,892 (4.6%; CI:[4.4%;4.8%]) | 16,906 (4.2%; CI:[3.9%;4.5%]) |

* Of the 13,011 FunFams with EC annotations considered in this analysis, 10% contained more than one EC annotation (*impure FunFams*). Applying DBSCAN resulted in 26,464 clusters and 74,706 (4.5%) sequences classified as outliers. On average, impure FunFams were split into more clusters than pure FunFams as indicated by the higher fold increase (3.9 compared to 1.8). The fold increase (number of clusters / number of FunFams) is given in brackets. CI indicates symmetric 95% confidence intervals.

**Fig. S4: Number of clusters for pure and impure FunFams.**

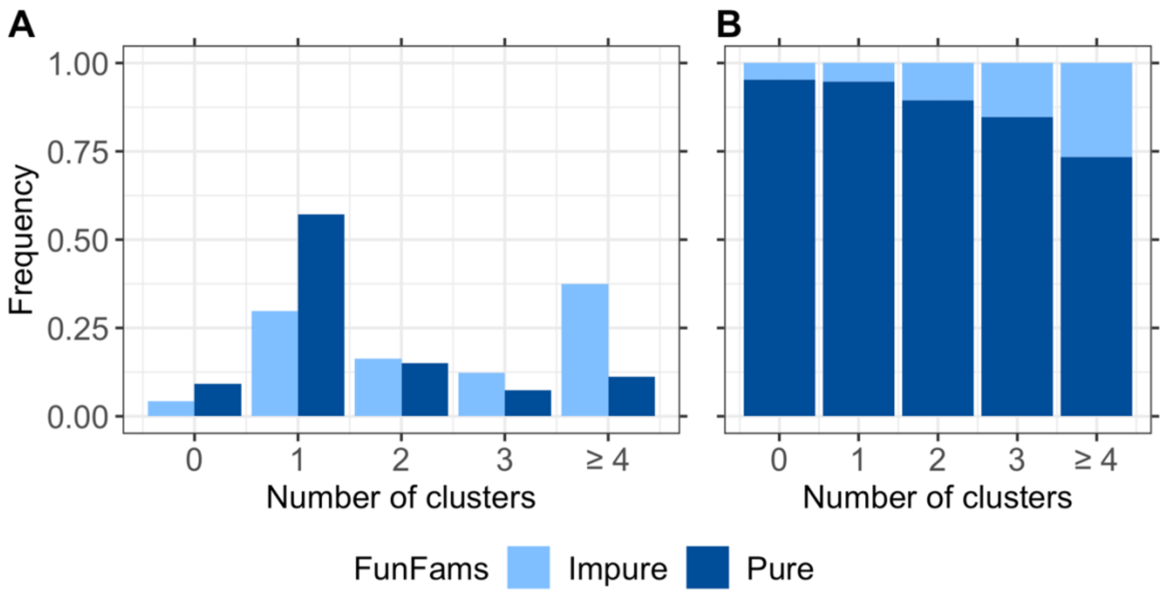

**A.** Impure FunFams are on average split into more clusters than pure ones. 66% of impure FunFams were split into at least two clusters while on 34% of pure FunFams were split. **B.** The fraction of impure FunFams increases with a higher cluster number, e.g., of all FunFams that were split into at least four clusters, 27% were impure while only 10% of all FunFams are impure. Bars at zero clusters (leftmost bars) indicate FunFams for which all sequences were classified as outliers.

##### 2.2. No consistent trend for assessment on different levels of EC annotations.

58% of all impure FunFams consisted of ECs which are identical on the first three and different on the fourth level (Fig. S5A), i.e., most of the FunFams were impure due to differences on the fourth level of EC annotations. For this level, it is also most difficult to distinguish sequences with different function. Two proteins with different first-level ECs are more likely evolutionarily unrelated, and therefore, less sequence-similar than two proteins identical in the first three EC levels.

Although a meaningful assumption, we failed to observe a clear trend, i.e., splitting FunFams into more consistent sub-families appeared neither harder nor easier if differences in EC annotations were on lower levels. The percentage of impure clusters was lowest for “level 2” (EC annotations were different on the second level) (Fig. S5C). The percentage of FunFams completely pure after clustering and the average purity were highest for “level 2” while the values were similar for the other levels (Fig. S5D&E). Those differences for level 2 could be mainly due to the small number of FunFams (Fig. S5A, 8%) with EC impurity on this level.

**Fig. S5: No consistent trend for assessment on different levels of EC annotations.**

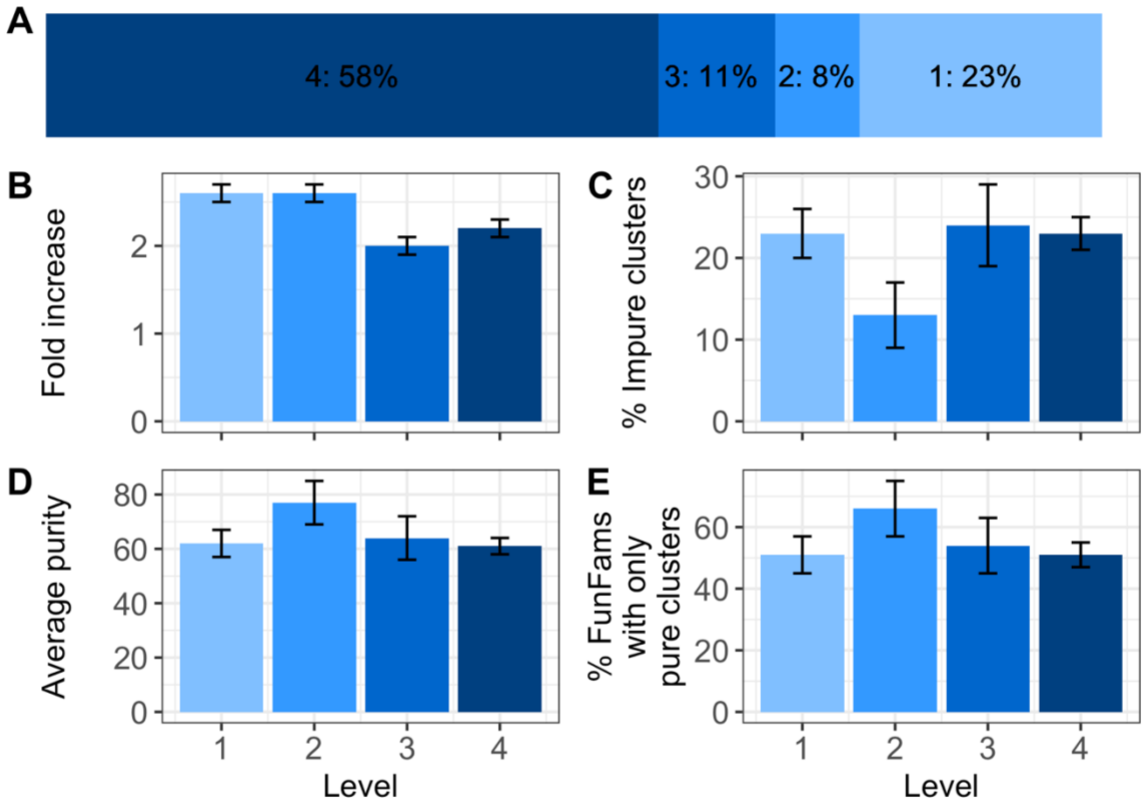

We split the set of impure FunFams into four subsets. Each subset for level *x* consists only of FunFams for which the EC annotations are identical until level *x-1* and different for level *x*. **A.** 58% of all impure FunFams are impure because of differences only at the fourth level of EC while still 23% of impure FunFams consist of sequences with EC annotations already different on the first level. **B.** The fold increase was similarly high for “level 1” and “level 2”, and lower for the other levels. Fold increase indicates the average number of clusters a FunFam was split into. **C.** The percentage of impure clusters was similarly high for “level 1” and “level 4”, lower for “level 3” and lowest for “level 2”. **D.** The percentage of completely pure FunFams after clustering was highest for “level 2” and lowest for “level 1”. **E.** The average purity increased for less specific EC levels (smaller numbers) for levels 2-4 while it was lowest for level 1.

### 3. More detailed assessment for five superfamilies

To assess the effect of the choice of embeddings and clustering parameters, and to allow a more detailed assessment of the clustering performance for different levels of EC annotations, we picked five different superfamilies with different properties (Table S5): (i) high number of moonlighting proteins, i.e., proteins with multiple EC numbers, (ii) a lot of divergence on the third level of EC numbers, i.e., the impure FunFams in this family contain a lot of annotations that are different in the third level of the EC numbers, (iii) a lot of divergence on the fourth level of EC numbers, i.e. the impure FunFams in this family contain many annotations that are identical until the third level, but are different on the fourth level, (iv) clustering worked well, i.e., the average purity per FunFam was high, and (v) clustering did not work well, i.e., the average purity per FunFam was low (Table S5).

**Table S5: Five interesting superfamilies chosen for more detailed analysis. ***

| Superfamily | Property | Number of FunFams | Number of impure FunFams |
|---|---|---|---|
| **3.40.50.150** (Vaccinia Virus protein VP39) | Number of moonlighting proteins | 302 | 28 |
| **3.20.20.70** (Aldolase class I) | Divergence in EC3 | 298 | 29 |
| **3.40.47.10** (Thiolase/Chalcone synthase) | Divergence in EC4 | 52 | 12 |
| **3.50.50.60** (FAD/NAD(P)-binding domain) | Good clustering | 161 | 17 |
| **1.10.630.10** (Cytochrome p450) | Bad clustering | 76 | 26 |

* To allow a more detailed analysis, we picked five exemplary superfamilies following five different criteria: (i) high number of moonlighting proteins, i.e. proteins with multiple EC numbers, (ii) a lot of divergence on the third level of EC numbers, i.e., the impure FunFams in this family contain a lot of annotations that are different in the third level of the EC numbers, (iii) a lot of divergence on the fourth level of EC numbers, i.e. the impure FunFams in this family contain many annotations that are identical until the third level, but are different on the fourth level, (iv) clustering worked well, i.e. the average purity per FunFam was high, and (v) clustering did not work well, i.e. the average purity per FunFam was low.

The fold increase, i.e., the number of clusters a FunFam is split into, was largest for superfamily 3.40.47.10 (with high divergence on the fourth level of EC annotations) and smallest for superfamily 1.10.630.10 (for which clustering worked badly) (Fig. S6A). The percentage of outliers was highest for 3.50.50.60 (for which clustering worked well) and lowest again for superfamily 1.10.630.10 (Fig. S6B). These numbers already indicated that the embedding distances between sequences in FunFams of superfamily 1.10.630.10 did not allow a differentiation between different functionalities of those sequences. Maybe the embeddings mainly captured certain structural constraints between sequences in this superfamily instead of zooming into the functional relations.

##### 3.1. Importance of parameter choice.

The distance threshold $\theta$ of DBSCAN (Ester *et al.*, 1996) defining whether two points are considered close to each other highly influences the clustering results. The variance observed between thresholds for different superfamilies indicated that it was reasonable to choose a threshold for each superfamily independently (Table S6).

**Table S6: Chosen distance cut-offs for clustering for five superfamilies. ***

| | **3.40.50.150** | **3.20.20.70** | **3.40.47.10** | **3.50.50.60** | **1.10.630.10** |
|---|---|---|---|---|---|
| **1st quartile** | 0.103 | 0.144 | 0.108 | 0.124 | 0.161 |
| **Median** | 0.176 | 0.238 | 0.203 | 0.202 | 0.209 |
| **3rd quartile** | 0.254 | 0.345 | 0.299 | 0.306 | 0.250 |

* The distance cutoff $\theta$ was chosen individually for each superfamily. For each sequence in a FunFam, we calculated the average distance of this sequence to all other sequences in this FunFam. From the resulting distribution, $\theta$ was chosen as the 1st quartile, median, or 3rd quartile.

Using smaller distances resulted in more clusters and outliers leading to a purer clustering (Table S7). Especially for superfamily 1.10.630.10, for which the default clustering did not work well (Table S5), using the 1st quartile of sequence distances as $\theta$ led to a much smaller number of impure clusters and a larger average purity than for the default clustering (Fig. S6). One major reason why the default clustering did not work well for superfamily 1.10.630.10 could be that the FunFams in this superfamily were mostly not split into any or only a small number of clusters (Fig. S6A; fold increase of 1.4, i.e., each FunFam was on average split into 1.5 clusters) and also only a low fraction of sequences was classified as outliers (Fig. S6B, 2% of sequences classified as outliers).

In addition to $n$=5, we tested fixed neighborhood sizes of $n \in [5;129;255]$ and variable neighborhood sizes dependent of the size of the FunFam $n$=x*|F| with |F|=number of sequences in a FunFam and x$\in$[0.01;0.1;0.2]. The individual performance differences between the five different superfamilies were not influenced by the neighborhood size, i.e., for the superfamily where the default clustering performed well/bad, also the clustering with any other neighborhood size, fixed or variable, performed well/bad (Fig. S6).

**Table S7: Influence of chosen parameters on resulting clustering. ***

| | | | Impure FunFams | | | |
|---|---|---|---|---|---|---|
| | **# Clusters** | **# Outliers** | **% Clusters with ECs** | **% Impure clusters** | **% FunFams with only pure clusters** | **Average purity** |
| **Default** | 1,603 | 3,760 | 61% (81%) | 13% (34%) | 50% (41%) | 59% (57%) |
| **ProtBERT** | 1,402 | 2,986 | 68% (86%) | 19% (41%) | 42% (37%) | 51% (49%) |
| **θ=1st quartile** | 2,261 | 8,221 | 53% (66%) | 4% (12%) | 73% (52%) | 83% (79%) |
| **θ=3rd quartile** | 1,144 | 1,544 | 72% (91%) | 30% (55%) | 29% (26%) | 37% (35%) |
| **n=129** | 423 | 11,464 | 98% (68%) | 35% (26%) | 60% (43%) | 60% (60%) |
| **n=255** | 325 | 6,960 | 94% (66%) | 38% (25%) | 57% (43%) | 57% (57%) |
| **n=0.05*\|F\|** | 1,211 | 8,087 | 72% (76%) | 21% (33%) | 52% (41%) | 58% (58%) |
| **n=0.1*\|F\|** | 1,034 | 11,644 | 84% (74%) | 27% (32%) | 54% (40%) | 58% (58%) |
| **n=0.2*\|F\|** | 851 | 17,249 | 91% (69%) | 35% (32%) | 56% (38%) | 57% (57%) |
| **Cosine** | 824 | 0 | 100% (100%) | 100% (100%) | 0% (0%) | 0% (0%) |
| **Manhattan** | 1,617 | 3,805 | 61% (81%) | 13% (34%) | 50% (41%) | 60% (57%) |
| **PIDE** | 2,074 | 10,464 | 45% (71%) | 12% (34%) | 42% (27%) | 57% (52%) |

* We show average clustering results for various parameters for five superfamilies (1.10.630.10, 3.20.20.70, 3.40.47.10, 3.40.50.150, 3.50.50.60). *n* indicates the chosen neighborhood size for DBSCAN, i.e., the number of sequences a sequence has to be close to be considered a core point. θ is the chosen distance cutoff to define whether a pair of sequences are close to each other. θ was chosen individually for each superfamily. Default: Clustering with same parameters as remaining analysis (*n*=5, θ=median of average distance of all sequences to all other sequences in the same FunFam for one superfamily); ProtBERT: ProtBERT embeddings (1024 dimensions) were used to represent sequences instead of PB-Tucker embeddings (128 dimensions, optimized to distinguish CATH classes); θ=1st quartile: *n*=5, θ=1st quartile of average sequence distances; θ=3rd quartile: *n*=5,

θ=3rd quartile of average sequence distances; *n*=x (x∈ [129,255]): *n*=x, θ=median of average sequence distance; n=x*|F| (x ∈ [0.05,0.1,0.2]): the neighborhood size was chosen individually for the each FunFam with *n*=max(x*|F|, 5) and |F|=number of sequences in FunFam F, θ=median of average sequence distance; Cosine: Cosine distance was used as distance measure; Manhattan: Manhattan distance was used as distance measure; PIDE: 1-PIDE was used as distance measure (PIDE: pairwise sequence identity).

**Fig. S6: Influence of chosen clustering parameters.**

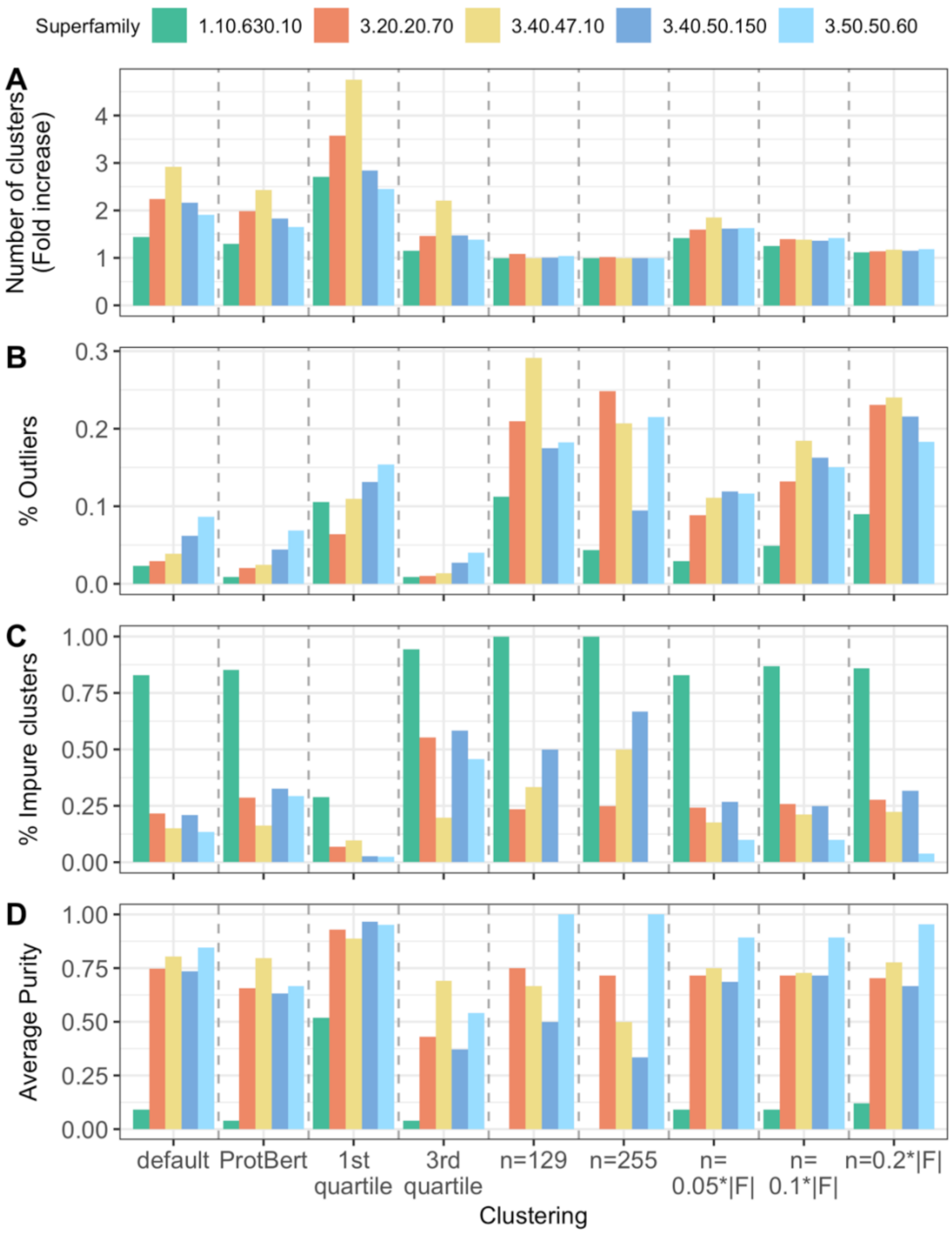

We show the individual clustering results for various parameters for five superfamilies (1.10.630.10, 3.20.20.70, 3.40.47.10, 3.40.50.150, 3.50.50.60). *n* indicates the chosen neighborhood size for DBSCAN, i.e., the number of sequences a sequence has to be close to be considered a core point. $\theta$ is the chosen distance cutoff to define whether a pair of sequences are close to each other. $\theta$ was chosen individually for each superfamily. Default: Clustering with same parameters as remaining analysis ($n$=5, $\theta$=median of average distance of all sequences to all other sequences in the same FunFam for one superfamily);

ProtBERT: ProtBERT embeddings (1024 dimensions) were used to represent sequences instead of PB-Tucker embeddings (128 dimensions, optimized to distinguish CATH classes); θ=1st quartile: *n*=5, θ=1st quartile of average sequence distances; θ=3rd quartile: *n*=5, θ=3rd quartile of average sequence distances; n=x (x∈[129,255]): *n*=x, θ=median of average sequence distance; n=x*|F| (x∈[0.05,0.1,0.2]): the neighborhood size was chosen individually for the each FunFam with *n*=max(x*|F|, 5) and |F|=number of sequences in FunFam F, θ=median of average sequence distance. **A**. The number of clusters as measured by the fold increase (i.e., increase over the number of FunFams) was especially high when using a small value for θ, e.g., here the 1st quartile of average sequence distances. There is no consistent relation between the number of clusters and the criterion for choosing a superfamily. **B**. The fraction of sequences classified as outliers exploded for larger values of n∈(129, 255) while it remained in a similar range for other parameter choices and independent of the criterion for choosing a superfamily. **C.** As expected, the fraction of impure clusters was very large for superfamily 1.10.630.10 which was chosen as example for "clustering did not work well", and for superfamily 3.50.50.60 chosen as example for "clustering worked well", the fraction of impure clusters was very low. **D.** The average purity of any of the five superfamilies was very similar for all choices of clustering parameters, except for θ=1st quartile where the average purity for superfamily 1.10.630.10 was much larger than for the other parameter sets. Therefore, if one is interested in having very pure clusters without being concerned about the large number of resulting clusters, using a smaller value for θ can be a reasonable approach.

##### 3.2. No consistent influence of level of difference in EC annotations.

Assessing how well our clustering approach worked depending on the level on which EC annotations were different (level *x*=ECs are identical until level x-1 and different on level *x*) did not reveal a consistent trend either for the full data set or the chosen five superfamilies.

For superfamily 3.40.47.10, we did not observe any impure FunFam for levels 1-3 because this family was chosen as family with high divergence only on the fourth level of EC, i.e., all proteins are annotated to the same EC class until the third level but are annotated to many different classes on the fourth level (Table S5). Therefore, 100% of all impure FunFams were impure due to differences in the EC annotations on the fourth level (Fig. S7).

For superfamily 1.10.630.10 for which clustering did not work well, the percentage of impure clusters was always highest, and consequently, the average purity was always lowest compared to the other four superfamilies (Fig. S7B&C) except for level 2 because no FunFam of this superfamily was impure due to differences on the second EC level. However, while the clustering led to almost no increase in purity for level 4, it worked better for level 1 and 3 (Fig. S7B&C, dark green bar). The percentage of FunFams impure on level 4 was lower than for the other superfamilies except for 3.20.20.70 which was specifically chosen because of the high divergence on level 3 and higher (Fig. S7A). This indicated that the superfamily 1.10.630.10 was less well split into FunFams and the functional impurity present in these was caused by coarse-grained functional differences as reflected by EC numbers different on higher levels.

For superfamily 3.40.50.150 with many moonlighting proteins, clustering was perfect for levels 1-3 (Fig. S7B&C, dark blue bar). Errors in the clustering when evaluating the fourth level might be in general caused by the presence of those moonlighting proteins and our rather conservative definition of purity: If one protein is annotated to two EC numbers and another protein in the same cluster is only annotated to one of those two, we consider this cluster impure. Since this superfamily contains many moonlighting proteins, this issue could highly influence the performance of the clustering approach. In fact, 88% of the moonlighting proteins in the impure FunFams are in FunFams with impurity on level 4 where clustering worked less well than for FunFams with impurity on level 1-3 with only a small fraction of moonlighting proteins.

The average purity dropped for level 3 and 2 compared to level 1 for superfamily 3.50.50.60 (Fig. S7C, lighter blue bar). On average, clustering worked well for this superfamily and apparently these good results were mainly caused by the perfect clustering on level 4. Our assumption was that it should be easier to detect more coarse-grained functional inconsistencies (i.e., different annotations on the first or second EC level), however for the superfamily where the clustering worked well, the opposite was the case.

The trend for superfamily 3.20.20.70 was the same as for the overall data set: The clustering worked best (i.e., perfectly) for levels 2 and 3, and it worked slightly worse for level 1 than for level 4 (Fig. S7B&C, orange bar). Investigating the impure FunFams on level 1 showed that the conservative definition of same EC annotation could explain the observed trend. Seven FunFams were impure on the first level. Of those, five were clustered perfectly into pure FunFams. The remaining two consisted of sequences where some of the sequences were annotated to EC number 5.3.1.1 and some were annotated to EC numbers 5.3.1.1 and 4.2.3.3. Our clustering approach did not cluster these FunFams further.

**Fig. S7: No consistent trend for assessment on different levels of EC annotations.**

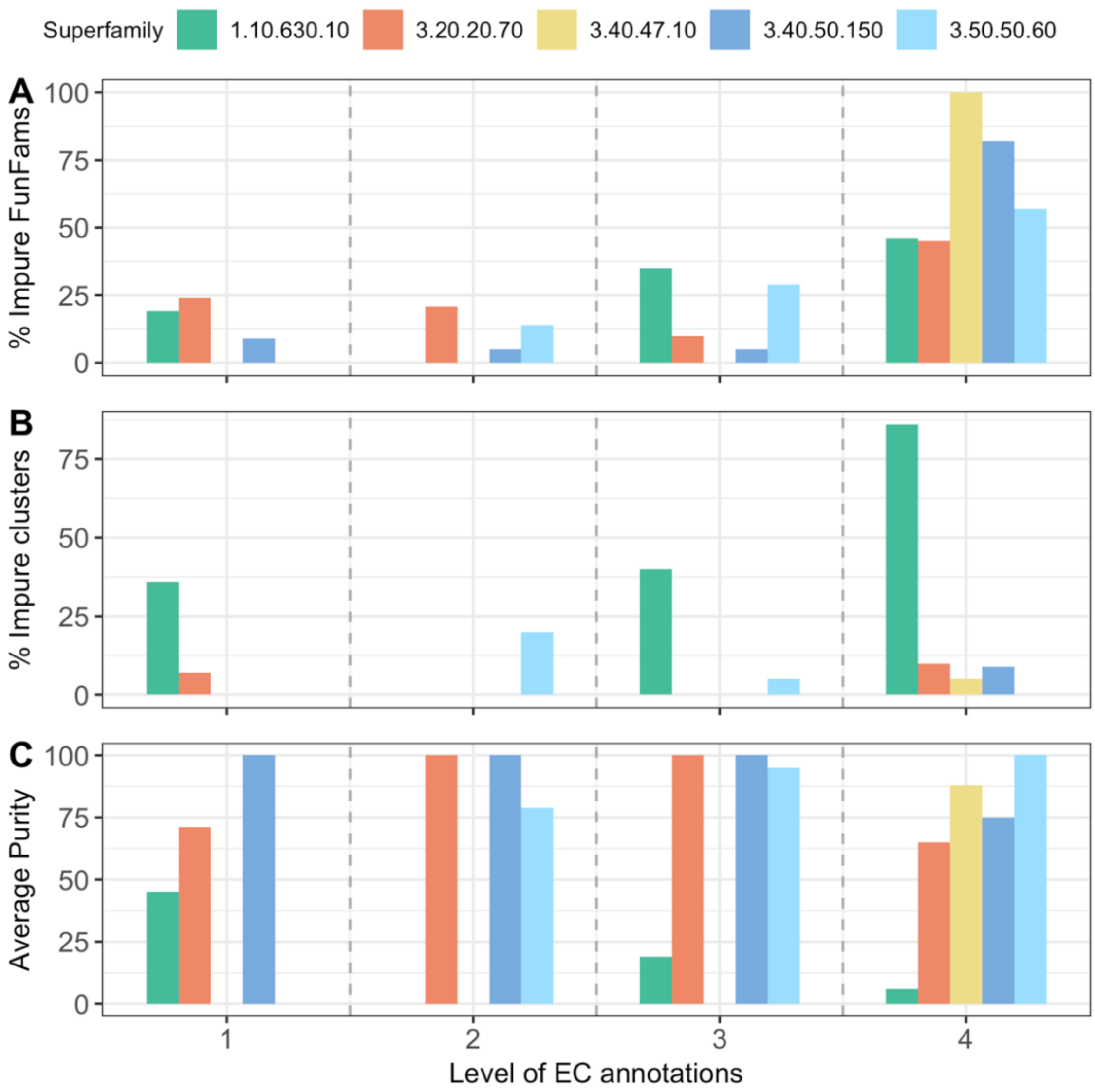

We assess purity of FunFams and clusters for five superfamilies (1.10.630.10, 3.20.20.70, 3.40.47.10, 3.40.50.150, 3.50.50.60) using different definitions of purity: A FunFam or cluster is considered "pure" on level x of EC annotations if the EC annotations of all sequences in that FunFam/cluster are identical at least until level x. On level 1, all annotations are considered identical that have the same first EC number, while on level 4, annotations are only considered identical if they are the same on all four levels of EC numbers. A. The fraction of impure FunFams per superfamily dropped for lower levels of EC annotations. For superfamily 3.40.47.10, we only observed impure FunFams for the fourth EC level because this family was particularly chosen that way (Table S2). Superfamily 1.10.630.10 for which clustering did not work well always had the highest fraction of impure FunFams on all levels of EC annotations. B. The fraction of impure clusters dropped for superfamily 1.10.630 while still remaining high, and clustering was perfect for superfamily 3.40.50.150 for levels 1-3. For the other two superfamilies, the fraction of impure clusters rose. Since the fraction of impure clusters was only calculated for impure FunFams (for a pure FunFams, all resulting clusters are pure), there were no impure clusters for superfamily 3.40.47.10 just because there were also no impure FunFams at levels 1-3. C. By definition, the opposite trends as observed for the fraction of impure clusters was true for the average purity. The drop in average purity for superfamilies 3.20.20.70 and 3.50.50.60 indicated that the FunFams which were not clustered correctly were mainly the FunFams containing EC annotations that were different even up to the first or second level.

### References for Supporting Online Material

Berman, H. M. *et al.* (2000) 'The Protein Data Bank', *Nucleic Acids Research*, 28(1), pp. 235–242. doi: 10.1093/nar/28.1.235.

*CATH: Protein Structure Classification Database at UCL* (2020). Available at: https://www.cathdb.info/ (Accessed: 2 November 2020).

Elnaggar, A. *et al.* (2020) 'ProtTrans: Towards Cracking the Language of Life's Code Through Self-Supervised Deep Learning and High Performance Computing', *bioRxiv*, p. 2020.07.12.199554. doi: 10.1101/2020.07.12.199554.

Ester, M. *et al.* (1996) 'A density-based algorithm for discovering clusters in large spatial databases with noise', in. *Kdd*, pp. 226–231.

*RCSB PDB* (2020). Available at: http://www.rcsb.org/ (Accessed: 20 November 2020).

Smyth, M. S. and Martin, J. H. J. (2000) 'x Ray crystallography', *Molecular Pathology*, 53(1), pp. 8–14. doi: 10.1136/mp.53.1.8.

Steinegger, M., Mirdita, M. and Söding, J. (2019) 'Protein-level assembly increases protein sequence recovery from metagenomic samples manyfold', *Nature Methods*, 16(7), pp. 603–606. doi: 10.1038/s41592-019-0437-4.

Steinegger, M. and Söding, J. (2017) 'MMseqs2 enables sensitive protein sequence searching for the analysis of massive data sets', *Nature Biotechnology*, 35(11), pp. 1026–1028. doi: 10.1038/nbt.3988.

Steinegger, M. and Söding, J. (2018) 'Clustering huge protein sequence sets in linear time', *Nature Communications*, 9(1), p. 2542. doi: 10.1038/s41467-018-04964-5.

Suzek, B. E. *et al.* (2015) 'UniRef clusters: a comprehensive and scalable alternative for improving sequence similarity searches', *Bioinformatics*, 31(6), pp. 926–932. doi: 10.1093/bioinformatics/btu739.

Tyzack, J. D. *et al.* (2018) 'Ranking Enzyme Structures in the PDB by Bound Ligand Similarity to Biological Substrates', *Structure*, 26(4), pp. 565-571.e3. doi: 10.1016/j.str.2018.02.009.

Yang, J., Roy, A. and Zhang, Y. (2013) 'BioLiP: a semi-manually curated database for biologically relevant ligand–protein interactions', *Nucleic Acids Research*, 41(Database issue), pp. D1096–D1103. doi: 10.1093/nar/gks966.
